# Red-light (670 nm) therapy reduces mechanical sensitivity and neuronal cell death, and alters glial responses following spinal cord injury in rats

**DOI:** 10.1101/2020.02.22.960641

**Authors:** Di Hu, Gila Moalem-Taylor, Jason R Potas

## Abstract

Individuals with spinal cord injury (SCI) often develop debilitating neuropathic pain, which may be driven by neuronal damage and neuroinflammation. We have previously demonstrated that treatment using 670 nm (red) light irradiation alters microglia/macrophage responses and alleviates mechanical hypersensitivity at 7-days post-injury. Here, we investigated the effect of red-light on the development of mechanical hypersensitivity, neuronal markers, and glial response in the subacute stage (days 1-7) following SCI. Wistar rats were subjected to a mild T10 hemi-contusion SCI or sham surgery followed by daily red-light treatment (30 min/day; 670 nm LED; 35mW/cm^2^) or sham treatment. Mechanical sensitivity of the rat dorsum was assessed from 1-day post-injury and repeated every second day. Spinal cords were collected at 1, 3, 5 and 7-days post-injury for analysis of myelination, neurofilament protein NF200 expression, neuronal cell death, reactive astrocytes (GFAP+ cells), interleukin1β (IL1β) expression, and inducible nitric oxide synthase (iNOS) production in IBA1^+^ microglia/macrophages. Red-light treatment significantly reduced the cumulative mechanical sensitivity and the hypersensitivity incidence following SCI. This effect was accompanied by significantly reduced neuronal cell death, reduced astrocyte activation and reduced iNOS expression in IBA1^+^ cells at the level of the injury. However, myelin and NF200 immunoreactivity and IL1β expression in GFAP^+^ and IBA1^+^ cells were not altered by red-light treatment. Thus, red-light therapy may represent a useful non-pharmacological approach for treating pain during the subacute period after SCI by decreasing neuronal loss and modulating the inflammatory glial response.

## Introduction

The World Health Organisation estimates between 250,000 and 500,000 people suffer from spinal cord injury (SCI) each year globally with an incidence of up to 1000 per million^1,2^. About 65-80% of spinal cord injured patients develop neuropathic pain^3–6^, which is considered severe in 20-30% of cases^7^. Multitherapeutic approaches utilising combinations of pharmacological intervention, exercise, massage and physiotherapy are often employed to manage neuropathic pain, however, these approaches require significant resources from different professionals and family members, and are generally inefficient^7,8^.

Neuropathic symptoms, including sensory hypersensitivity (e.g. stimulus-evoked pain, dysesthesia or pinprick hyperalgesia) in dermatomes corresponding to the lesion level are common is patients with SCI^9^. Numerous mechanisms have been implicated in the development of neuropathic pain following SCI, including structural damage (e.g. apoptosis, demyelination, cytoskeletal damage), biochemical damage and excitotoxicity, neuroinflammation (e.g. glial activation, cytokine secretion), enzyme dysregulation, and neuronal hyperexcitability^10^. Microglia/macrophages, astrocytes, and oligodendrocytes can significantly influence neuropathic pain post SCI^11^. For example, a major proinflammatory cytokine secreted by glial cells, interleukin-1 beta (IL1β), is believed to sensitize dorsal horn neurons that are responsible for both the development of, and providing positive feedback to enhance and maintain neuropathic pain^12–16^. Nitric oxide (NO), which is mainly derived from microglial inducible NO synthase (iNOS), has been shown to modulate long-term potentiation of C-fibres that eventually leads to hypersensitivity^17^. Several studies have shown that by altering some of these key inflammatory mediators, it might be possible to stall the development of neuropathic pain, or even depress its severity following SCI^18–21^.

Light therapy refers to the application of light irradiation, using laser or light emitting diodes (LED), to modulate the cellular biology in the pursuit of clinical benefits. Wavelengths between 630 nm and 830 nm have been demonstrated to reduce glial activation and reduce secretion of inflammatory cytokines in different injury models, including SCI^22,23^ We have previously established that 670 nm treatment in spinal cord injured rats reduces microglia/macrophage activation and alleviates mechanical hypersensitivity at 7-days post-injury (dpi)^21^. In the present study, we examined the effects of 670 nm treatment during the subacute phase of recovery (prior to 7-dpi) following SCI on: i) the development of mechanical hypersensitivity; ii) myelination and neuronal apoptosis; and iii) astrocyte and microglia/macrophages presence in the spinal cord and their associated IL1β and/or iNOS expression in the white matter tracts.

## Methods

### Animals and Spinal cord injury

Sibling male Wistar rats, 52 ± 4 days old, were used for this study with ethics approvals from the ANU Animal Experimentation Ethics Committee. Animals were held in individually ventilated cages with environmental enrichment, standard food and water *ad libitum*. The housing facility was maintained at 20 °C with a 12-hour dark/light cycle. Animals were housed for 2 weeks prior to experimentation, and all experiments were carried out during the light cycle. Animals were anaesthetised using isoflurane (1.7-2.3 v/v %) while being maintained at 37 □ by a heat mat during all surgical procedures. Laminectomy was performed at the T10 vertebra where the dura mater and arachnoid were removed. A 10 g weight drop from 25 mm above the spinal cord was used to induce a hemi-contusion on the right side of the spinal cord^21,24^. Sham-injured animals underwent the same surgical procedures without the impaction. All animals received subcutaneous injections of antibiotics (Cephalothin Sodium; DBL) after surgery and once every 24 h throughout the recovery period at a dosage of 15 mg/kg/12 h. Animals were returned to their home cage after the effects of anaesthesia had subsided.

### Treatment

Spinal cord injured animals were divided into untreated (SCI; n=40) and 670 nm treated (SCI+670; n=33) groups. Sham-injured animals were divided into untreated (sSCI; n=12) and 670 nm treated (sSCI+670; n=11) groups. Treatments commenced 2 h after the surgery and were repeated every 24 h after behavioural sensitivity testing. For the duration of the treatment, animals were contained in a transparent Perspex box in their own cages. A commercially available LED array (75 mm^2^) that provided light at 670 ± 15 nm with a measured irradiance of 35.4 ± 0.05 mW/cm^2^ (WARP 75A; Quantum Devices, Inc) was placed directly above a Perspex box for light-treated animals (SCI+670 and sSCI+670 groups). With this light source, we have previously measured the irradiance at the ventral surface of the spinal cord to be 3.2 ± 0.6 mW/cm^2^ in a similar cohort of animals, as well as have described details of the spectral features and irradiance measurements of this light source^21^. The treatments were delivered for 30 mins per day (63.7 ± 0.09 J/cm^2^ per session) to the dorsal surface of the animal. Untreated animals (SCI and sSCI groups) were handled in the same way without the LED being turned on (sham-treatment).

### Sensitivity testing

All animals were subjected to sensitivity testing on every odd day from 1-day post injury (dpi) as we have previously described^21^. All sessions were carried out by the same assessor, who was blinded to experimental groups. Briefly, a nylon filament (OD: 1.22 mm, mass delivered: 2.86 ± 0.09 g) was used to deliver non-noxious tactile stimuli to 6 defined regions: dermatomes innervated by nerves above the level of the injury (dermatomes C6-T3), at the level of the injury (dermatomes T9-T12) and below the level of the injury (dermatomes L2-L5) on both ipsilateral and contralateral sides. At each region, 10 consecutive stimuli were applied, and responses were categorized into four different categories: I) no response; II) acknowledgement of the stimulus; III) sign of pain avoidance behaviour including moving away from the stimulus; IV) severe pain avoidance behaviour including jumping, running and vocalisation. The categories were then assigned a weight individually, 0, 1, √2, 2 and the summation of the 10 responses gave the regional sensitivity score (RSS). The summation of the six RSSs gave the cumulative sensitivity score (CSS). A group of age-matched male Wistar rats were also included in the sensitivity testing as non-injured control animals (CON; n=7). A hypersensitivity threshold was defined as 2 standard deviations above the mean of CON animals. Hypersensitivity incidence was thus calculated as the percentage of animals whose CSS exceeded hypersensitivity threshold. See Hu *et al.,* 2016 for more details on sensitivity testing^21^.

### Tissue collection and processing

At the end of designated recovery periods (1, 3, 5 and 7-days post injury), animals were euthanized and perfused transcardially with 0.9% (w/v) saline followed by 4% (w/v) paraformaldehyde. The spinal cord was dissected, cryoprotected in 30% (w/v) sucrose solution for at least 24 h before cryosectioning. Spinal cord sections 1.5 mm rostral and 1.5 mm caudal to the injury epicentre were placed in Tissue-Tek OCT compound (IA018; Proscitech) and then sectioned horizontally at 20 μm thickness on a cryostat (CM1850; Leica Microsystems) at −20 °C.

### Terminal deoxynucleotidyl transferase dUTP nick end labelling

The same protocol was used as published before^21^. Slides were incubated with 1:10 TdT buffer for 10 min and subsequently in reaction mixture [0.12% v/v TdT (3333574001, Roche applied science); 0.25% v/v dUTP (11093070910, Roche applied science); 10.51% v/v TdT buffer] for 1 hr at 37 °C. This was followed by 15 min incubation with 1:10 SSC buffer and blocking with 10% v/v normal goat serum in 0.1M PBS for 10 min. Slides were then incubated with 0.1% v/v streptavidin 488 (S-11223, Invitrogen) at 37 °C for 30 min. This was followed by immunohistochemistry where appropriate.

### Immunohistochemistry

Standard immunohistochemistry procedures were followed. Slices around the injury epicentre were dehydrated in 70% ethanol and then rehydrated in distilled water and subsequently in 0.1M PBS. The antigen retrieval step involved incubation in Reveal-it solution (AR2002, ImmunoSolution) for 6-12 h at 37 □ followed by 0.1M PBS washes. Slides were blocked in 20% (v/v) normal donkey serum in 0.1M PBS with 0.1% *(w/v)* bovine serum albumin (BSA) for 1 hr at room temperature before the addition of primary antibodies. Primary antibodies [myelin basic protein (1:1000, Abcam ab40390); NeuN (1:200, Abcam ab104224); NF200 (1:1000, Invitrogen 131200); GFAP (1:1000, Dako Z0334); IL1β (1:200, R&D systems AF501-NA); IBA1 (1:200, Wako 019-19741 and Abcam ab5076); UNOS (1:150, Invitrogen PA1-039)] were diluted in 0.1M PBS containing 2% *(v/v)* normal donkey serum and 0.1% *(w/v)* BSA and incubated overnight at 4 □. Negative controls were also included where the primary antibodies were excluded. Slides were washed with 0.1M PBS followed by secondary antibody incubations at room temperature for 1-2 h. Secondary antibodies [goat anti mouse Alexa Fluor 488 (1:1000, Invitrogen A-11001); goat anti rabbit Alexa Fluor 594 (1:1000, Invitrogen A-31631); donkey anti-goat Alexa Fluor 594 (1:500, Abcam ab150132); donkey anti-rabbit Alexa Fluor 488 (1:1000, Invitrogen A-21206)] were diluted in the same way as primary antibodies and followed by 0.1M PBS washes. Slides were then incubated in 1:1000 *(v/v)* diluted Hoechst solution (94403; Sigma-Aldrich) and then washed off using 0.1 M PBS.

### Image acquisition and quantification

Images were scanned and imaged using a Nikon A1 confocal microscope fitted with camera (DS-Qi1; Nikon). In MBP/NF200/Hoechst staining, the entire cross section of the spinal cord at the injury epicentre was scanned using three lasers (405 nm, 488 nm and 561 nm) under x10 magnification with a z-plane of at least 10 μm in depth. In NeuN/TUNEL/Hoechst staining, 6 representative images were imaged in the dorsal, intermediate and ventral regions of the spinal cord on both ipsilateral and contralateral sides under x20 magnification using the same lasers and the same z-plane configurations as above. All imaging and laser settings were kept constant for all animals for each staining. Images were then transferred offline for analysis using ImageJ v1.46r^25^. NeuN/Hoechst and NeuN/TUNEL/Hoechst staining was analysed using Cell Counter plugin as described earlier^21^. NE200 was expressed as particles (density/mm^2^). The regions of interest were defined and quantified prior to cell counting covering a minimum area of 0.1 mm^2^. MBP was analysed as % area of fluorescence.

In GFAP/Hoechst staining, the entire cross-section of the spinal cord at the injury level was scanned using two lasers (405 nm and 488 nm) under ×10 magnification with a z-plane of at least 10 μm in depth. In GFAP/IL1β/Hoechst, IBA1/IL1β/Hoechst, and IBA1/UNOS/Hoechst staining, six representative images were taken in the dorsal, lateral and ventral spinal cord funiculi at the injury level on both ipsilateral and contralateral sides under ×20 magnification using three lasers (405 nm, 488 nm and 561 nm) and the same z-plane configuration as above. All imaging and laser settings were kept constant for all animals for each staining. Images were then analysed offline using ImageJ v1.46r^25^. GEAP was analysed as fluorescence per % area while GFAP/IL1β/Hoechst, IBA1/IL1β/Hoechst, and IBA1/UNOS/Hoechst were analysed using Cell Counter plugin as described earlier^21^. The areas of interest were defined and quantified prior to cell counting covering a minimum area of 0.1 mm^2^. Cell quantification is expressed as the number of cells per unit area (mm^2^).

### Statistical analysis

All data were expressed as mean ± SEM unless otherwise stated. Statistical analysis was carried out using R^26^. General linear mixed-effects models (GLMER; *glmer* function in R for binomially distributed data) or linear mixed-effects models implementing Satterthwaite’s approximation (LMER; *lmer* function in R) were used for multiple factor analysis accounting for random effects^27^ LMER was followed by Tukey or False Discover Rate post-hoc adjustments for pairwise comparisons implemented by estimated marginal means (*emmeans* fuction in R) where appropriate. Log transformations (log[x+1]) were performed when the data was heteroscedastic and improved model fitting. Model fitting was assessed by inspection of residual plots and calculating Akaike Information Criterion (*AIC* function in R) 28.

For data that failed to fit LMER models, a Cumulative Link Mixed Model was fitted, accommodating one random factor, with the Laplace approximation (CLMM; *clmm2* function in R)^29^. CLMM required categorising scores into bins that best represented the data. Histograms of complete data sets (blind to groups) were inspected to determine natural breaks in the data to define each category. CLMM models were assessed with Akaike Information Criterion as well as inspection by plotting the observed data against the model predicated values, and comparing a linear model made with these data, to a line with an intercept of zero and a slope of 1. Post-hoc pairwise comparisons were performed using a Wilcoxon rank sum test on raw scores where applicable, with Bonferroni p value correction.

Both models accommodated up to 4 fixed factors (treatment, side, regions and time), and one (CLMM: animal identification) or two (LMER: animal identification nested within their respective families) random factors. Models of CSSs were confirmed against models of RSSs, which enable more factors and thereby improved model fitting and statistical power. P values less than 0.05 were considered as statistically significant.

## Results

### Red-light reduces the level and incidence of mechanical hypersensitivity

To determine whether red-light treatment affects mechanical hypersensitivity during the subacute phase of SCI, we measured mechanical sensitivity in control, untreated and light-treated spinal cord-injured and sham-operated rats. We first assessed the range of cumulative sensitivity scores (CSSs) of uninjured control animals by quantifying the sum of regional sensitivity scores (RSSs) of uninjured normal control animals (CON; n=7; **Fig 1a).** Most animals had a mix of category I and category II responses, except for one animal that demonstrated two category III responses at the contralateral Above-Level region. The CON group had a mean CSS of 2.75 and a standard deviation of 2.09 and consequently a hypersensitivity threshold of 6.92 (mean + 2×SD) was established. The Above-Level region was more sensitive than the At-Level region (p = 0.041, LMER, Tukey post-hoc), while differences between sides failed to reach significance (p = 0.073, LMER).

**Fig 1.**
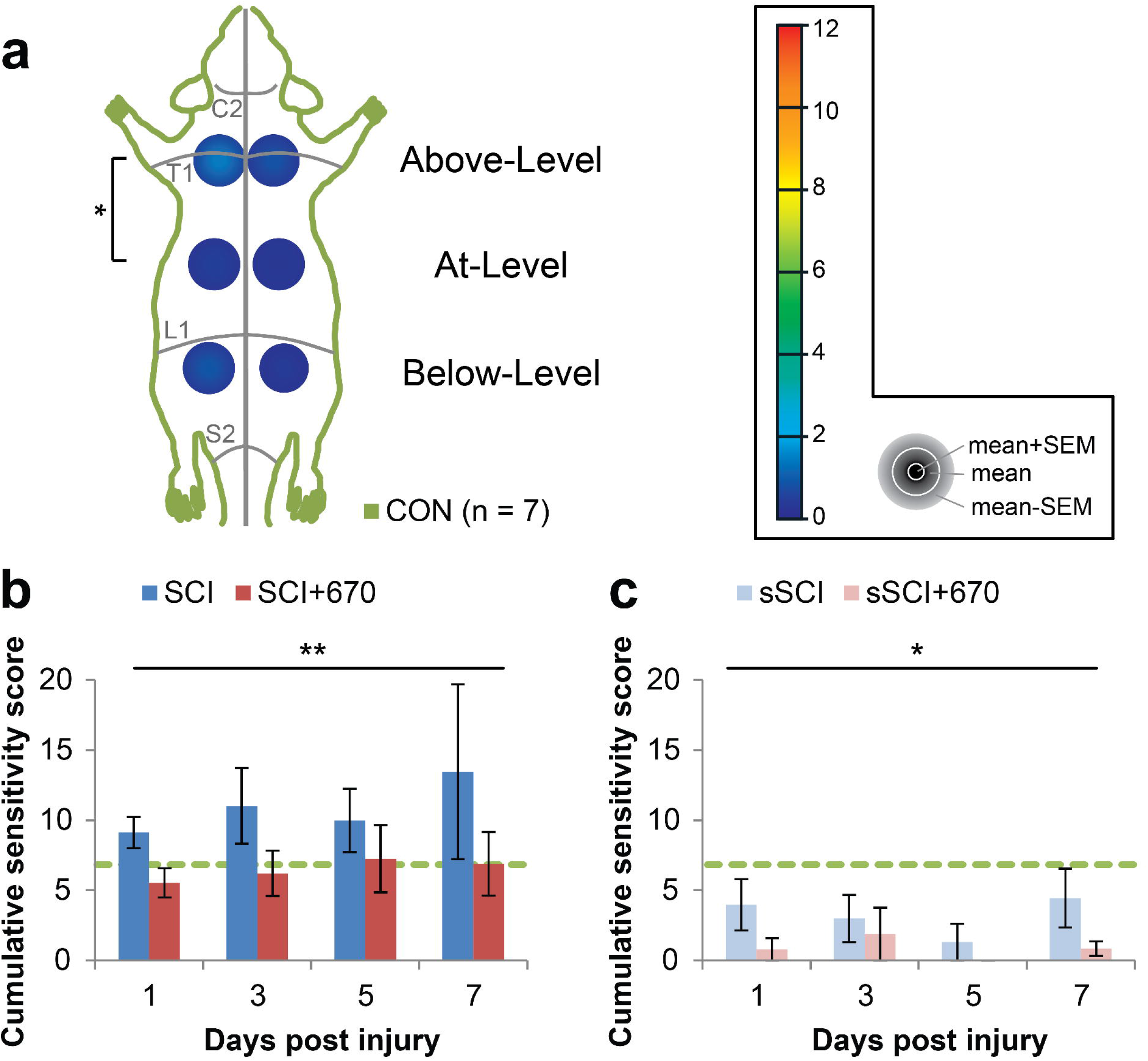
Cumulative sensitivity is increased up to 7-dpi following mild T10 hemi-contusion SCI but reduced after red-light treatment. **(a)** RSSs of the CON group (n = 7) showing the mean + SEM, mean, and mean — SEM in concentric order as indicated by the colour scale and accompanying grey legend (insert). RSSs, obtained from six regions (left and right sides; “Above-Level”, “At-Level”, and “Below-Level” relative to T10 injury), are overlayed on a schematic representation of the rat dorsum, with C2, T1, L1 and S1 dermatomes and midline indicated (grey lines). Statistical comparison between levels is indicated by the black bracket. **(b)** CSSs in all spinal cord injured animals with (SCI+670) or without (SCI) red-light treatment. **(c)** CSSs in all sham-injured animals with (sSCI+670) or without (sSCI) red-light treatment. Green-dashed line indicates hypersensitivity threshold (6.92) derived from CON group (see Methods). Statistical comparison between levels in CON group (vertical bracket, CLMM) and between SCI and SCI+670 groups across the time points (black line, CLMM) are indicated. Data is expressed as mean ± SEM; * p < 0.05, ** p < 0.01. See Fig 2 for n values for all groups at each time point. Abbreviations: CSS, cumulative sensitivity score (sum of RSSs); CON, non-injured control; RSS, regional sensitivity score; SCI, spinal cord injured untreated; SCI+670, spinal cord injured + red-light treatment; sSCI, sham-injured untreated; sSCI+670, sham-injured + red-light treatment.

We then quantified mechanical sensitivity responses at 1-7 days post-injury (dpi) in animals with SCI (SCI, untreated) and SCI with 670 nm light treatment (SCI+670, **Fig 1b;** see **Fig 2** for n values). Compared to the CON group, CSSs were significantly elevated in the SCI group (p = 0.040, LMER, Tukey), but not the SCI+670 group (p = 0.55, LMER, Tukey) over the 7-day recovery period. Furthermore, the SCI+670 group displayed a significant reduction of CSSs compared to the SCI group (p = 0.0097, LMER, Tukey). There was no time effect (p = 0.86) or interaction between treatment and time (p = 0.80) in the spinal cord injured groups **(Fig 1b).**

**Fig 2.**
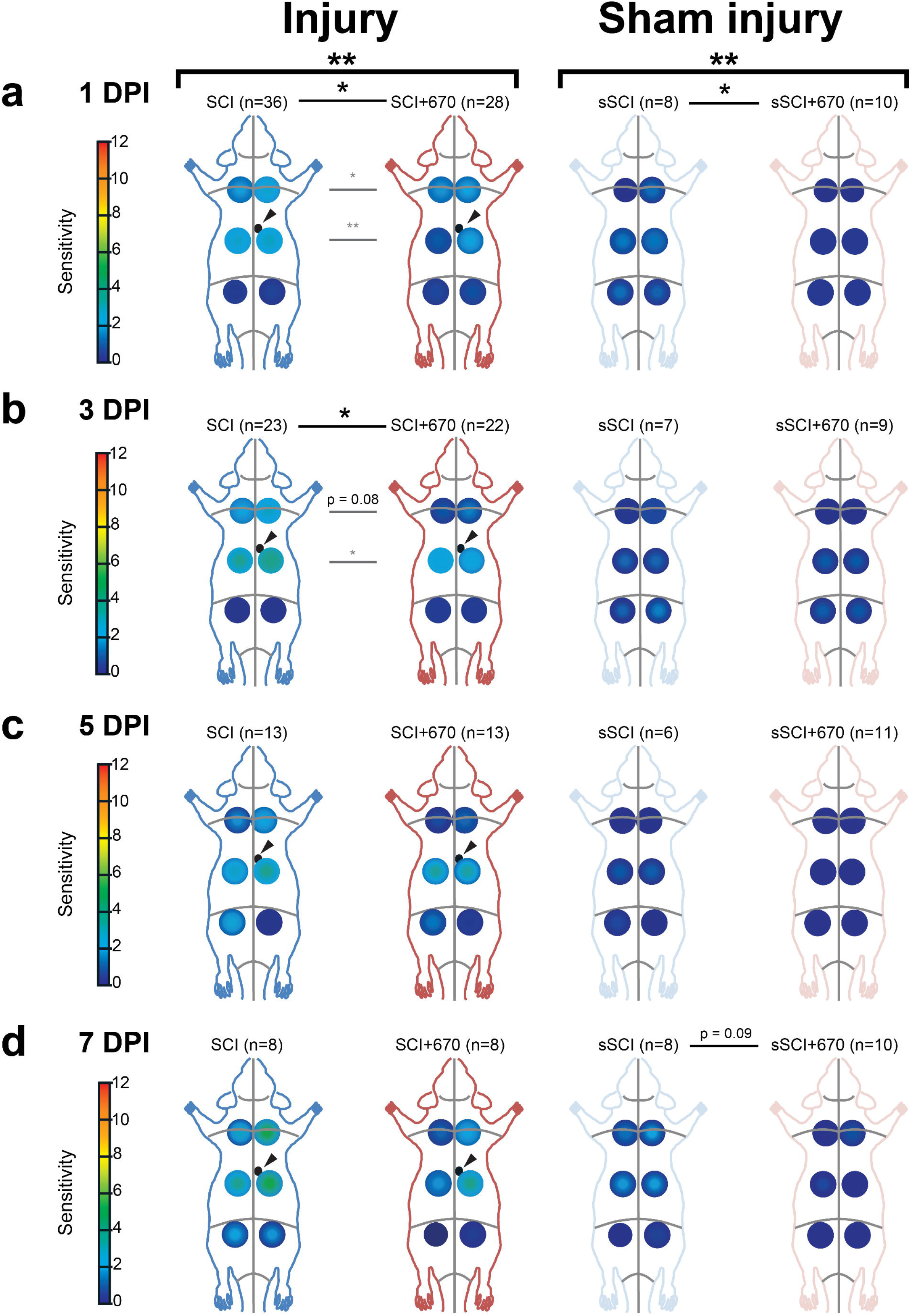
At-Level and Above-Level regional sensitivity are reduced by red-light treatment. RSSs in SCI (blue), SCI+670 (red), sSCI (light blue), and sSCI+670 (pink) animals at **(a)** 1-dpi, **(b)** 3-dpi, **(c)** 5-dpi, and **(d)** 7-dpi are shown for all animals investigated. Arrowheads indicate location of T10 hemi-contusion injury (small back circles) in spinal cord injured groups. Statistical comparisons between 2 groups across all time points (black bracket, CLMM), across all levels at individual time point (black lines, CLMM) and between 2 groups at different levels (grey lines, CLMM) are indicated. Data is expressed as mean ± SEM as per **Fig 1a** inset; n values indicated for each group; * p < 0.05, ** p < 0.01, p value indicated for 0.1 < p < 0.05. See **Fig 1** for abbreviations.

In sham-injured groups (sham-injured + untreated, sSCI; sham-injured + 670 nm treated, sSCI+670) that were subjected to the same sensitivity testing **(Fig 1c),** a significant effect of red-light treatment on CSSs was observed (p = 0.018, CLMM). Post-hoc analysis revealed that CCSs of sSCI animals were not significantly different to the CON group (p = 1.0, Wilcoxon rank sum, Bonferroni adjusted), however, interestingly, the sSCI+670 group were significantly reduced compared to both CON and sSCI groups (p = 0.0003 and p = 0.0075 respectively, Wilcoxon rank sum, Bonferroni adjusted).

To assess the spatial effects of injury and treatment on mechanical hypersensitivity, regional sensitivity scores (RSSs) in 6 regions were analysed across up to 7-dpi **(Fig 2, Sup 1&2).** Congruent with the analysis above, the SCI+670 group displayed an overall reduction in mechanical sensitivity compared to the SCI group (p = 0.004, CLMM; **Fig 2a-d,** left panel), and similarly, the sSCI+670 group also displayed a significant reduction in mechanical sensitivity compared to the sSCI group (p = 0.006; CLMM; **Fig 2a-d,** right panel). For injured groups, the treatment effect was most evident at 1-dpi (p = 0.010, **Fig 2a,** left) and 3-dpi (p = 0.024, **Fig 2b,** left), whereas for the sham-injured groups, the effect of red-light was most obvious at 1-dpi (p = 0.029, **Fig 2a,** right). Interestingly, a significant reduction of RSSs following red-light treatment in both injured and sham-injured groups was detected in animals that did not develop hypersensitivity **(Sup 1**).

To determine whether red-light treatment also affects the incidence of animals developing mechanical hypersensitivity (see **Sup 2** for RSSs), we analysed the proportion of animals with CSS > hypersensitivity threshold in all treatment groups from day 1 to day 7 following SCI or sham injury **(Table 1**). Following injury, over half of the SCI group developed hypersensitivity up to 5-dpi, and only approximately 1/3 in the SCI+670 group. While the effect of red-light significantly reduced the incidence of developing hypersensitivity over the 1 to 5-dpi period (p = 0.037, GLMER), the effect failed to reach significance across the entire 7-day period (p = 0.080, GLMER). There was no significant difference in the incidence of sham-injured animals reaching the hypersensitivity threshold between the two groups, however only a few animals in each group developed hypersensitivity **(Table 1)**.

**Table 1.** Red-light treatment reduces hypersensitivity incidence following a mild T10 hemicontusion SCI. Hypersensitivity incidence following SCI or sham-SCI is shown for untreated and red-light treated groups, as the percentage of animals with scores above the hypersensitivity threshold (see **Fig 1** for threshold, see **Sup 2** for RSSs of hypersensitive animals).

| Days post injury | SCI<br>(% hypersensitive) | SCI+670<br>(% hypersensitive) | sSCI<br>(% hypersensitive) | sSCI+670<br>(% hypersensitive) |
| --- | --- | --- | --- | --- |
| 1-dpi | 56% (n = 20) | 32% (n = 9) | 25% (n = 2) | 10% (n = 1) |
| 3-dpi | 57% (n = 13) | 31% (n = 7) | 14% (n = 1) | 11% (n = 1) |
| 5-dpi | 54% (n = 7) | 23% (n = 3) | 17% (n = 1) | 0 |
| 7-dpi | 38% (n = 3) | 50% (n = 4) | 25% (n = 2) | 0 |

These behavioural results indicate that red-light treatment reduces mechanical sensitivity scores following both SCI and sham injury, and that red-light treatment reduces the incidence of developing hypersensitivity for at least the first 5 days.

### Red-light does not affect the level of myelination and heavy neurofilament expression

To determine whether red-light treatment affects axons in the injured spinal cord, immunohistochemical analysis was carried out to assess the degree of myelination (MBP staining) and density of axons (NF200 expression) at different time points post-injury.

The percentage area of positive MBP staining was analysed from both sides over the recovery points in both spinal cord injury groups **(Fig 3).** On the side contralateral to the injury **(Fig 3c),** the percentage area positive for MBP staining was around 50-60% throughout the recovery period in both groups. This level of staining was similar to the CON group **(Fig 3c).** On the ipsilateral side of the spinal cord **(Fig 3d),** there was significantly less area of MBP^+^ staining compared to the contralateral side in both injury groups (p = 1.5e-08, LMER). Compared to the CON group, the ipsilateral side was significantly reduced in both the SCI (p = 0.004, LMER) and SCI+670 (p = 0.0362, LMER) groups.

**Fig 3.**
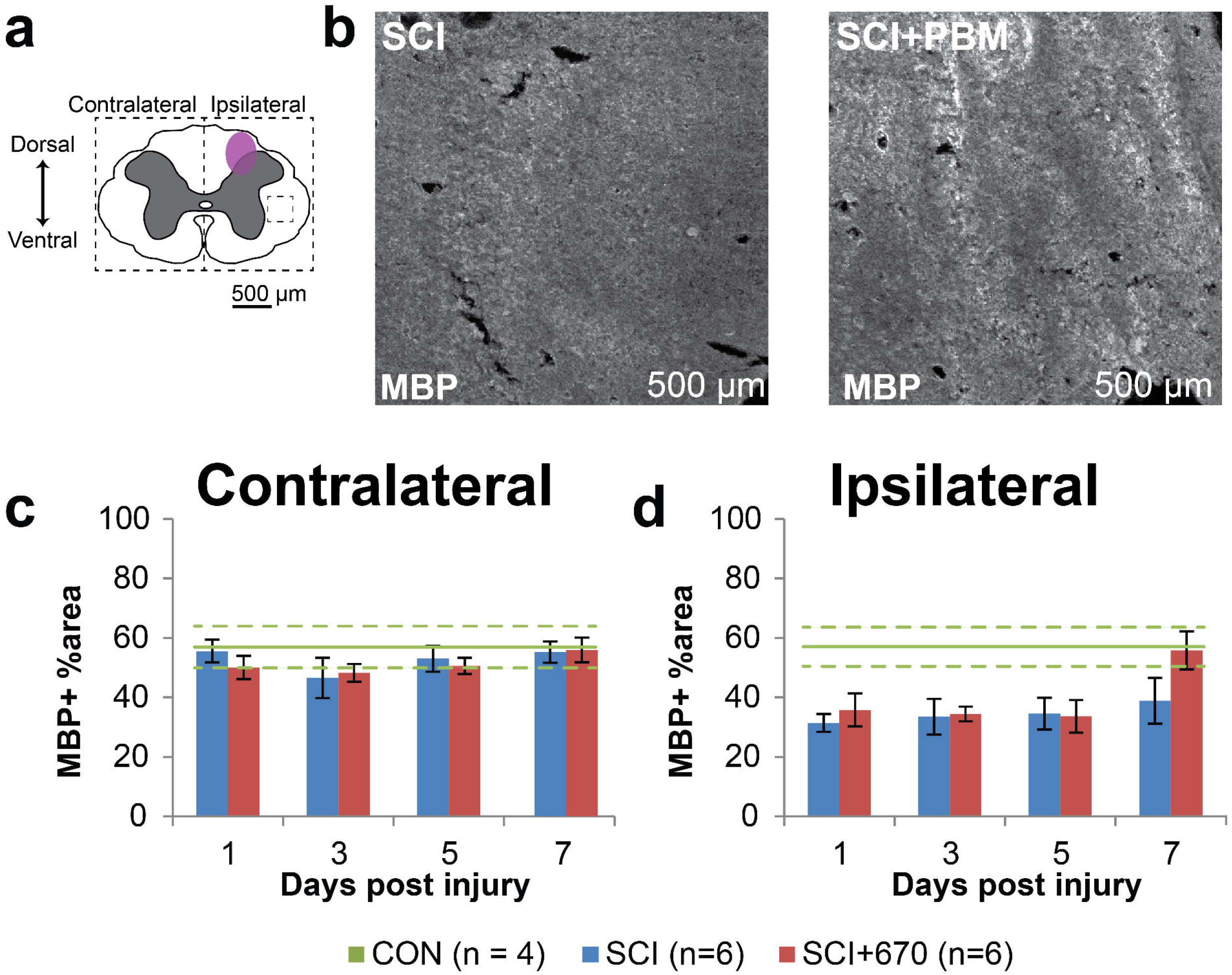
Ipsilateral demyelination was observed from 1-5 dpi post hemi-contusion SCI. **(a)** Schematic representation of the spinal cord illustrates the approximate location of the injury epicentre (purple shaded area) and region of interest (dotted) for MBP labelling quantification. **(b)** Example images are shown of positive MBP labelling (white) from sham- and light-treated groups ipsilateral to the injury at the dorsal level at 7-dpi (region of example images indicated by small dashed box in **a), (c-d)** Quantification of MBP positive labelling, expressed as the percentage area of positive label above background within the region of interest (dotted boundaries in **a)** contralateral **(c)** and ipsilateral **(d)** to the injury of sham- and light-treated groups. Data for control animals are shown (solid green, mean; dotted green, SEM All other data expressed as mean ± SEM; n values indicated (legend) are for each time point.

Analysis of the heavy neurofilament NE200 in the spinal cord grey matter following injury with and without 670 nm treatment **(Fig 4)** was carried out from three regions (dorsal, intermediate, and ventral) across both sides of the spinal cord **(Fig 4a).** In CON animals, all six regions showed comparable levels of NE200 density (a total average of 12575 ± 915 particles/mm^2^) with no significant difference on either side (p = 0.94, LMER), region (p = 0.50, LMER), or combinations of side and region (p = 0.97, LMER) **(Fig 4c-h).** Following SCI, there was an overall significant reduction in NF200 density across all regions and sides of the spinal cord compared to the control group (SCI, p = 0.0002; SCI+670, p = 0.0004; LMER, Tukey). No significant differences between the SCI and SCI+670 groups across all regions and sides was observed (p = 0.71, LMER). The dorsal regions showed a significant reduction compared to the intermediate (p = 0.031, LMER, Tukey) and ventral (p = 0.0002, LMER, Tukey) regions. Across all regions and sides, the 7-dpi showed an overall significant reduction in NF200 density compared to earlier time points (1-3 dpi; p ≤ 0.037, LMER, Tukey). This time effect arose from the intermediate (p = 0.0012; LMER, Tukey) and ventral (p = 5.6e-6, LMER, Tukey) regions **(Fig e-h),** but not the dorsal region (p = 0.79, LMER, Tukey, **Fig c-d),** which displayed a significant reduction of NF200 particle density on the ipsilateral side (p = 0.014, LMER, Tukey).

**Fig 4.**
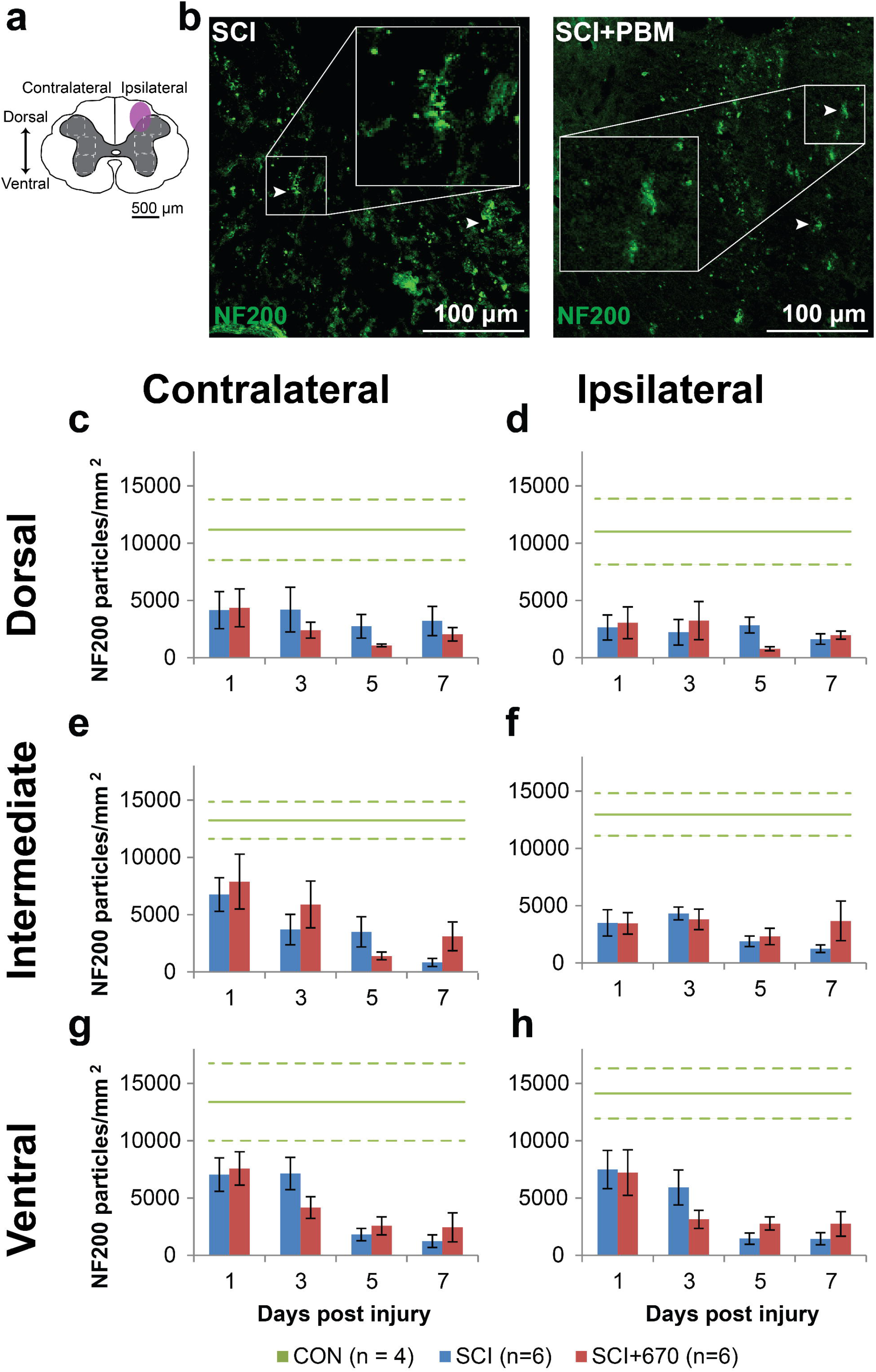
NF200 density is reduced following SCI but not altered by red-light treatment. **(a)** The schematic representation of the spinal cord illustrates the dorsal, intermediate and ventral regions of interest for analysis (enclosed by dashed lines, 0.1 mm^2^). Approximate location of injury is indicated (purple shaded area). **(b)** Example images of NF200 (green) positive labelling from spinal cord injured sham- and light-treated groups ipsilateral to the injury at dorsal level at 7-dpi. **(c-d)** Quantification of NF200+ labelling expressed as positive particle density within the region of interest, in the dorsal region of the spinal cord, contralateral **(c)** and ipsilateral **(d)** to the injury of sham- and light-treated groups. **(e-f)** NF200+ particle density in the intermediate regions of interest contralateral **(e)** and ipsilateral **(f)** to the injury, **(g-h)** NF200+ particle density in the ventral regions of interest contralateral **(g)** and ipsilateral **(h)** to the injury. Data for control animals are shown (solid green, mean; dotted green, SEM). All other data expressed as mean ± SEM; n values indicated for each time point (legend). See text for statistical comparisons.

### Red-light ameliorates the level of neuronal cell death

We stained the spinal cords against NeuN/DAPI **(Sup 3),** as well as NeuN/TUNEL/DAPI which marks neuronal cells undergoing apoptosis or necrosis **(Fig 5).** There was no significant difference between groups in NeuN cell density, despite a strong reduction in the ipsilateral side across all levels (p < 2e-16, LMER) **(Sup 3).** Analysis of neuronal cell death was carried out on the six regions of interest across both sides of the spinal cord grey matter **(Fig 5a).** Examples of NeuN/TUNEL/DAPI staining in both SCI and SCI+670 groups are shown in **Fig 5b.** The vast majority of triple labelled cells arose at 1-dpi across all sides and regions **(Fig 5c-h).** At this time point, red-light treatment significantly reduced the density of NeuN^+^TUNEL^+^DAPI^+^ cells across all regions from the ipsilateral (p = 0.0009, LMER), but not contralateral side (p = 0.28, LMER).

**Fig 5.**
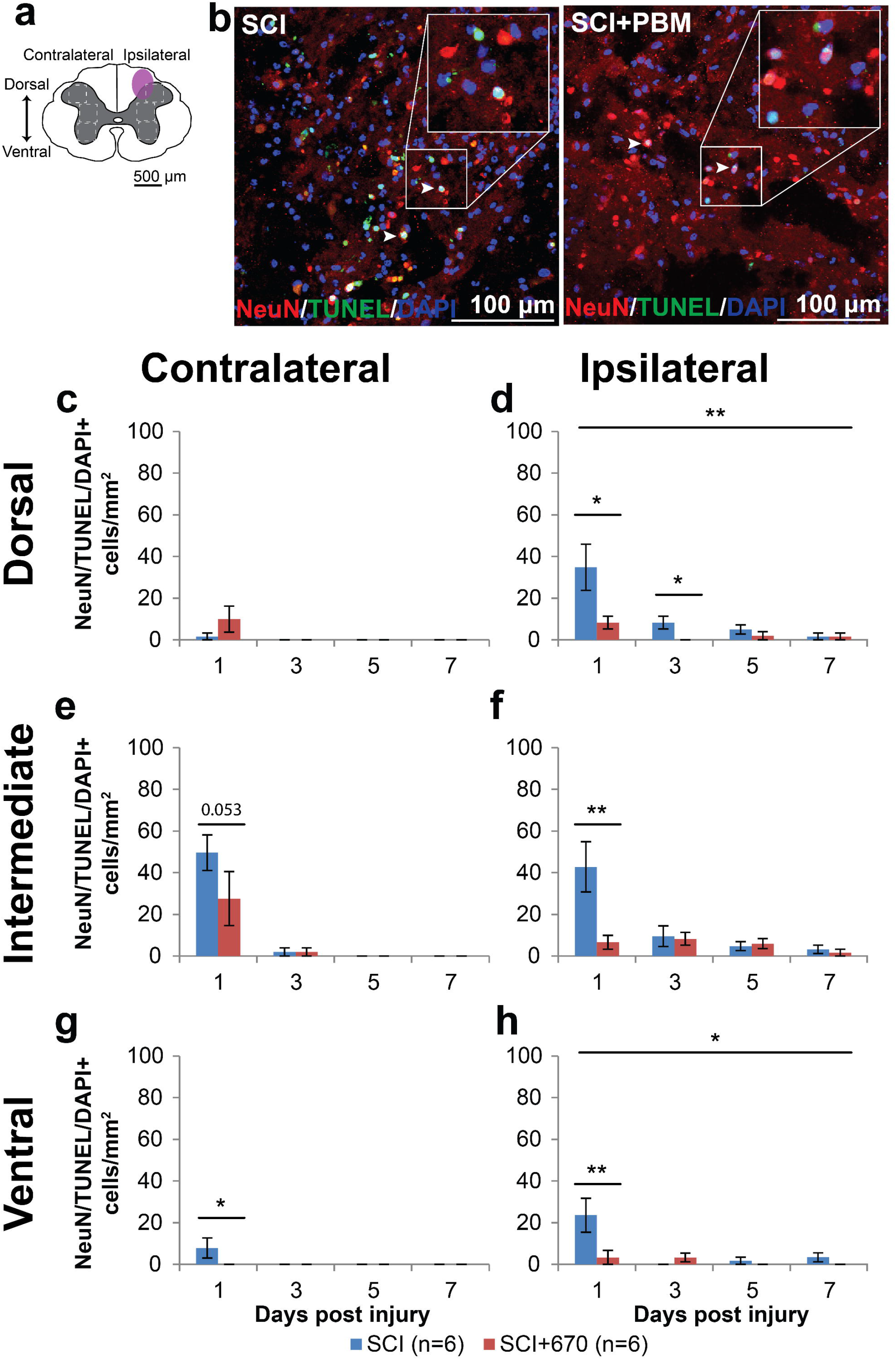
Neuronal cell death occurs early following SCI which is reduced by 670 nm treatment. **(a)** The schematic representation of the spinal cord illustrates the dorsal, intermediate and ventral regions of interest for analysis (enclosed by dashed lines, 0.1 mm^2^). Approximate location of injury epicentre is indicated by the purple shaded area. **(b)** Example images of NeuN (red), TUNEL (green), and DAPI (blue) triple positive cells from spinal cord injured untreated and light-treated groups ipsilateral to the injury at dorsal region at 1-dpi. **(c-d)** Quantification of NeuN^+^TUNEL^+^DAPI^+^ cells, expressed as triple positive cell density within the region of interest, in the dorsal region of the spinal cord, contralateral **(c)** and ipsilateral **(d)** to the injury of untreated and light-treated groups. **(e-f)** NeuN^+^TUNEL^+^DAPI^+^ cell density in the intermediate regions of interest contralateral **(e)** and ipsilateral **(f)** to the injury. **(g-h)** NeuN^+^TUNEL^+^DAPI^+^ cell density in the ventral regions of interest contralateral **(g)** and ipsilateral **(h)** to the injury. Data is expressed as mean ± SEM; n values indicated (legend) are for each time point. Statistical comparison between SCI and SCI+670 (LMER) are shown; * p < 0.05, ** p < 0.01.

### Red-light reduces astrocyte reactivity in the spinal cord

We have previously demonstrated reductions in activated microglia/macrophages following 670 nm treatment in spinal cord injured rats after 7 days of recovery^21^. To examine whether red-light treatment also influences astrogliosis in the injured spinal cord, astrocyte activation was quantified as the percentage area of GFAP^+^ immunofluorescence across 6 regions of the spinal cord **(Fig 6a).** Examples of GFAP^+^ staining for SCI and SCI+670 groups are shown in **Fig 6b.** Across groups, spinal cord regions and time, GFAP+ area was significantly increased ipsilateral to the injury compared to the contralateral side (p = 6.3e-14, LMER). Across all spinal cord regions, sides and time, red-light significantly reduced the GFAP^+^ area (p = 0.0081, LMER), which mainly arose from 3- to 7-dpi (p = 0.0063, LMER). A group difference was evident in the dorsal **(Fig 6c-d,** p = 0.0027, LMER) and ventral **(Fig 6g-h,** p = 0.046, LMER) regions of the spinal cord over the 3-7 day recovery period, but this failed to reach significance in the lateral region **(Fig 6e-f,** p = 0.148, LMER).

**Fig 6.**
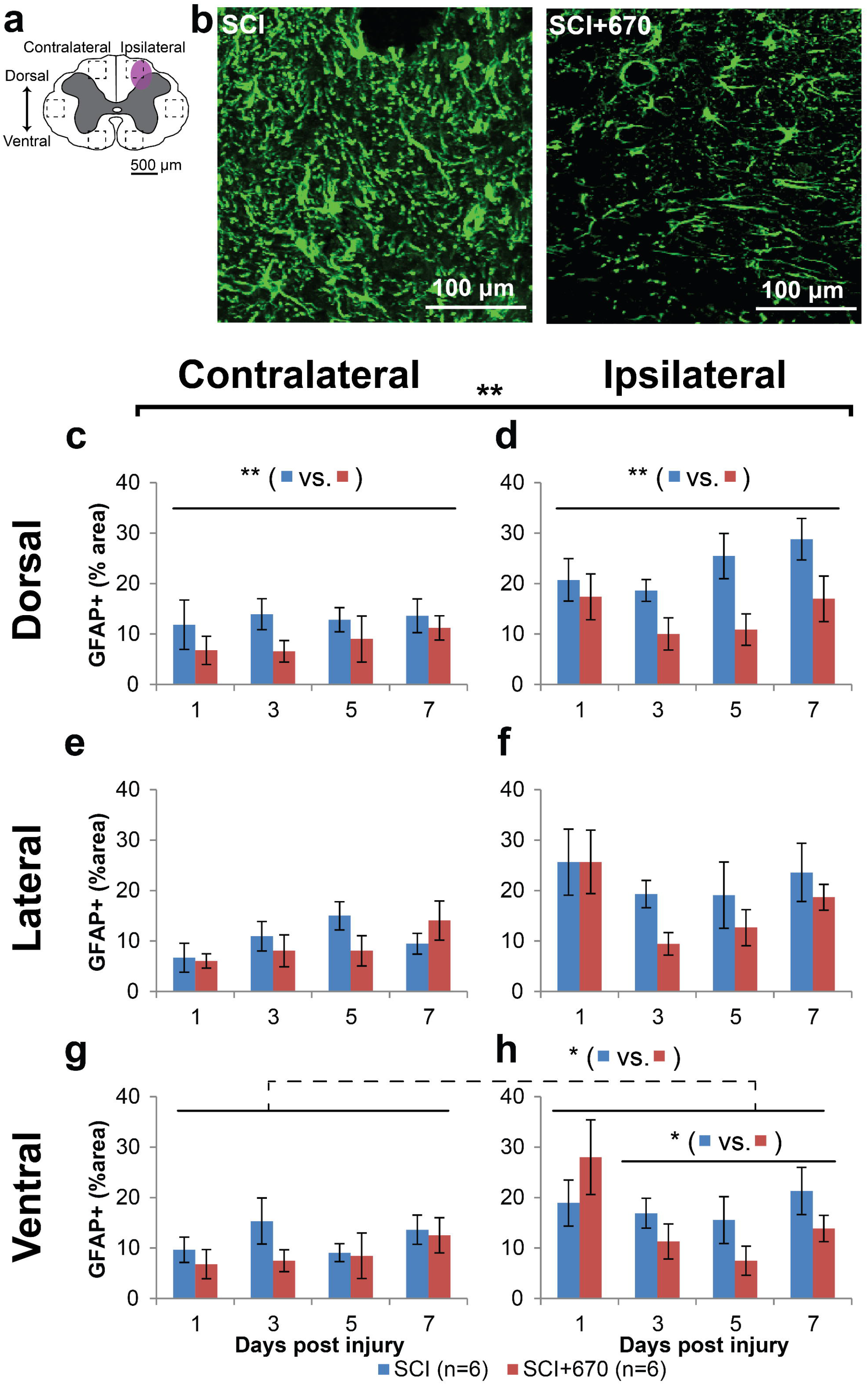
SCI-induced astrocyte activation is reduced following red-light treatment. **(a)** The schematic representation of the spinal cord illustrates the dorsal, lateral and ventral regions of interest for analysis (enclosed by dashed lines, area of each box: 0.1 mm^2^). Approximate location of injury is indicated by the purple shaded area. **(b)** Example images are shown of positive GFAP labelling (green) from untreated and light-treated groups ipsilateral to the injury at the dorsal region at 3-dpi. **(c-d)** Quantification of GFAP^+^ labelling, expressed as the percentage area of positive label above threshold within the dorsal regions of interest contralateral **(c)** and ipsilateral **(d)** to the injury of untreated and light-treated groups. **(e-f)** GFAP^+^ label in the lateral regions of interest contralateral **(e)** and ipsilateral **(f)** to the injury, **(g-h)** GFAP+ label in the ventral regions of interest contralateral **(g)** and ipsilateral **(h)** to the injury. n values indicated (legend) are for each time point. Statistical comparisons between SCI and SCI+670 groups across all time points and regions (black bracket) and across the time points at different region (black line) are indicated. Dotted indicates significance across both sides. Data is expressed as mean ± SEM; * p < 0.05, ** p < 0.01 (LMER). See **Fig 1** for abbreviations.

To determine if GFAP levels on the contralateral side at 1-dpi were elevated compared to control animals, GFAP expression 24 hours post-injury (SCI, n = 4) and sham-injury (sSCI, n = 4) were compared to normal animals (CON, n = 4; **Sup 4).** Overall, 24 hours after injury, GFAP^+^ area was elevated in the SCI group on both sides of the spinal cord compared to the control group (contralateral, p = 0.001, ipsilateral p < 0.001, LMER, Tukey) and sSCI animals (contralateral, p = 0.007, ipsilateral p < 0.001, LMER, Tukey). Compared to the control animals, the sSCI animals failed to reach significant difference on the contralateral side (p = 0.60, LMER, Tukey), and was significantly increased but with a small effect size in the ipsilateral side (p < 0.049. LMER, Tukey; **Sup 4c).**

These results demonstrate that astrocytes are already elevated across both sides of the spinal cord following hemi-contusion as early as 1-dpi at the spinal segment of injury, but more so on the ipsilateral side. GFAP expression is not affected by red-light at 1-dpi, but is significantly supressed by 670 nm treatment from 3-dpi onwards.

### Red-light does not affect IL1β expression in glial cells

IL1β is a pro-inflammatory cytokine that plays a key role in pain signalling^30^. We therefore investigated IL1β expression in astrocytes and microglia/ macrophages in 6 regions across the spinal cord **(Fig 7a; Fig 8a)** following hemi-contusion and 670 nm treatment. Examples of IL1β^+^GEAP^+^ cells, defined as triple positive for IL1β, GEAP and DAPI, are shown in **Fig 7b,** and of IBA1, IL1β and DAPI, are shown in **Fig 8b.**

**Fig 7.**
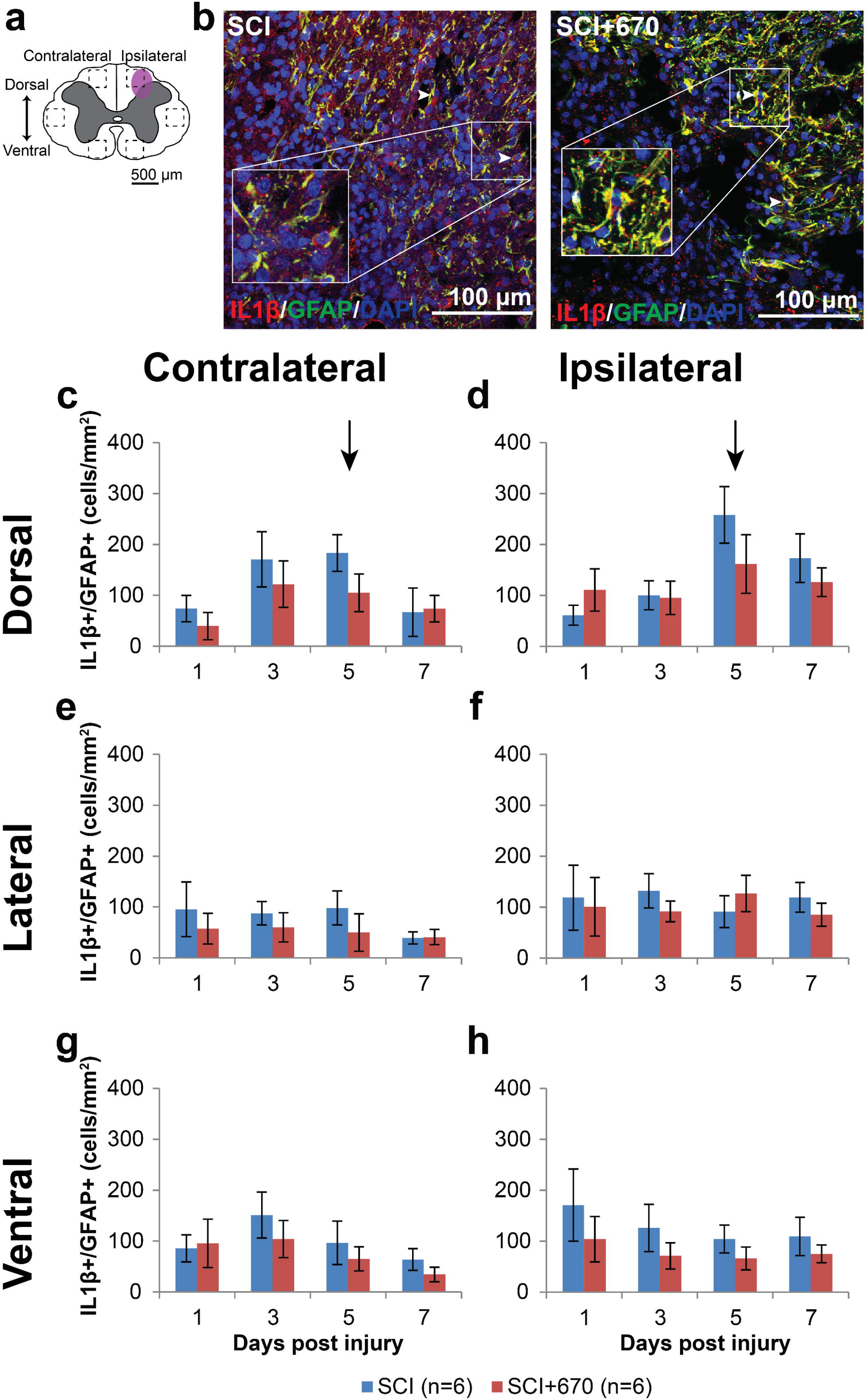
The density of IL1β producing astrocytes is not affected by red-light treatment following mild T10 hemi-contusion. **(a)** The schematic representation of the spinal cord illustrates the dorsal, lateral and ventral regions of interest for analysis (enclosed by dashed lines, area of each box: 0.1 mm^2^). Approximate location of injury is indicated by the purple shaded area. **(b)** Example images of IL1β (red), GEAP (green), and DAPI (blue) triple positive cells from spinal cord injured untreated and light-treated groups ipsilateral to the injury at dorsal region at 7-dpi. **(c-d)** Quantification of IL1β^+^GEAP^+^DAPI^+^ cells, expressed as triple positive cell density within the region of interest, in the dorsal region of the spinal cord, contralateral **(c)** and ipsilateral **(d)** to the spinal cord injury of untreated and light-treated groups. **(e-f)** IL1β^+^GEAP^+^DAPI^+^ cell density in the lateral regions of interest contralateral **(e)** and ipsilateral **(f)** to the injury, **(g-h)** IL1β^+^GEAP^+^DAPI^+^ cell density in the ventral regions of interest contralateral **(g)** and ipsilateral **(h)** to the injury. n values indicated (legend) are for each time point. Arrows indicate the 5-dpi time point is significantly increased in the dorsal region compared to lateral and ventral regions (p = 0.0003, LMER, Tukey). Data is expressed as mean ± SEM See **Fig 1** for abbreviations.

Throughout the spinal cord **(Fig 7c-h),** IL1β^+^GEAP^+^ cell density was elevated on the ipsilateral side compared to the contralateral side (p = 2.7e-07, LMER), while there was no overall significant effect of red-light treatment (p = 0.18). At the dorsal region, a time effect was apparent (p = 0.046, LMER), notably that the 5-dpi timepoint was significantly elevated compared to the 1-dpi timepoint across both sides (p = 0.030, LMER, Tukey). This significant IL1β^+^GEAP^+^ cell density elevation at the 5-dpi time point in the dorsal region (arrows, **Fig 7c-d)** was also elevated compared to the same time point at the lateral **(Fig 7e-f)** and ventral **(Fig 7g-h)** regions (p = 0.0003, LMER, Tukey).

IL1β^+^IBA1^+^ cell density was mostly under 50 cells/mm^2^ across the spinal cord regions, which was significantly reduced compare to IL1β expressing astrocytes (p = 1.7e-35, paired t-test; compare **Fig 7** with **Fig 8).** Throughout the 6 regions **(Fig 8c-g),** the ipsilateral side to the injury showed consistently increased IL1β^+^IBA1^+^ cell density compared to the contralateral side (p = 1.2e-5, LMER). Red-light treatment had no significant effect on IL1β^+^IBA1^+^ cell density (p = 0.97, LMER).

**Fig 8.**
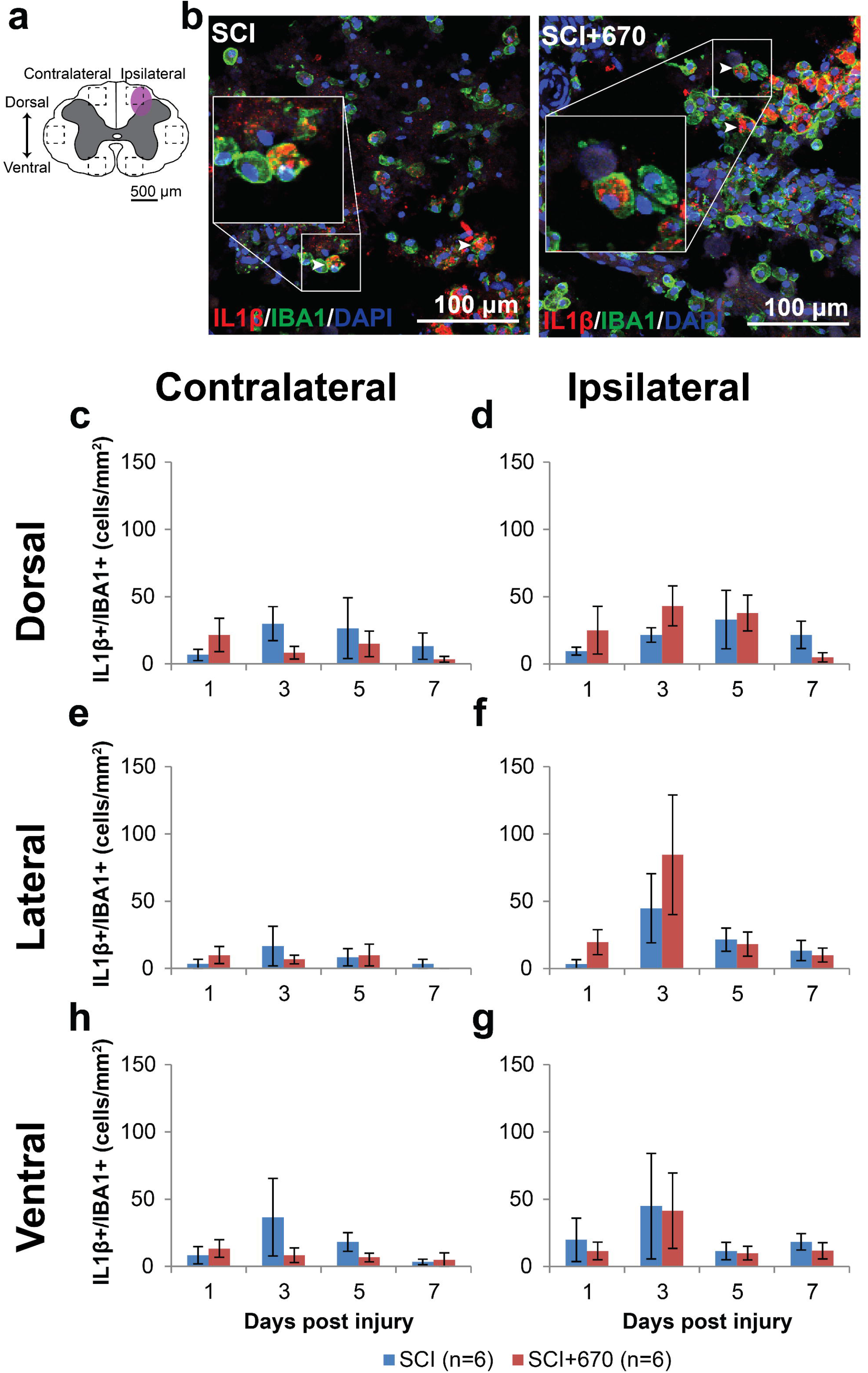
IL1β producing microglia/macrophage population is not affected by red-light treatment following T10 hemi-contusion. **(a)** The schematic representation of the spinal cord illustrates the dorsal, lateral and ventral regions of interest for analysis (enclosed by dashed lines, area of each box: 0.1 mm^2^). Approximate location of injury is indicated by the purple shaded area. **(b)** Example images shown of IL1β (red), IBA1 (green), and DAPI (blue) triple positive cells from spinal cord injured untreated and light-treated groups at the dorsal level, ipsilateral to the injury at 7-dpi. **(c-d)** Quantification of IL1β^+^IBA1^+^DAPI+ cells, expressed as triple positive cell density within the region of interest, are shown for the dorsal region of the spinal cord, contralateral **(c)** and ipsilateral **(d)** to the injury of untreated and light-treated groups. **(e-f)** IL1β^+^IBA1^+^DAPI^+^ cell density in the lateral regions of interest contralateral **(e)** and ipsilateral **(f)** to the injury. **(g-h)** IL1β^+^IBA1^+^DAPI^+^ cell density in the ventral regions of interest contralateral **(g)** and ipsilateral **(h)** to the injury. n values indicated (legend) are for each time point. Data is expressed as mean ± SEM See **Fig 1** for abbreviations.

These results demonstrate that IL1β-expressing glial cells were present throughout the spinal cord following hemi-contusion with higher numbers on the ipsilateral side. Red-light treatment had no significant effect on IL1β expression from either astrocytes or microglia/macrophages.

### Red-light reduces iNOS producing microglia/macrophages at the injury zone

NO is produced by iNOS in microglia/macrophages, and contributes to neuropathic pain processing^31^. Microglia/macrophages produce the iNOS isoform that can be detected by UNOS staining, which labels all three nitric oxide synthase isoforms (eNOS, nNOS, and iNOS). Six spinal cord regions **(Fig 9a)** were triple stained for UNOS/IBA1/DAPI to investigate the density of iNOS producing microglia/macrophages in the spinal cord following injury and red-light treatment. An example of staining is shown in **Fig 9b.** Throughout the 6 regions **(Fig 9c-h),** there was significantly greater UNOS+/IBA1+ cell density on the ipsilateral side (p = 1.2e-05, LMER), a strong effect of time (p = 0.0005), and a significant reduction in the red-light treated group (p = 0.0203).

**Fig 9.**
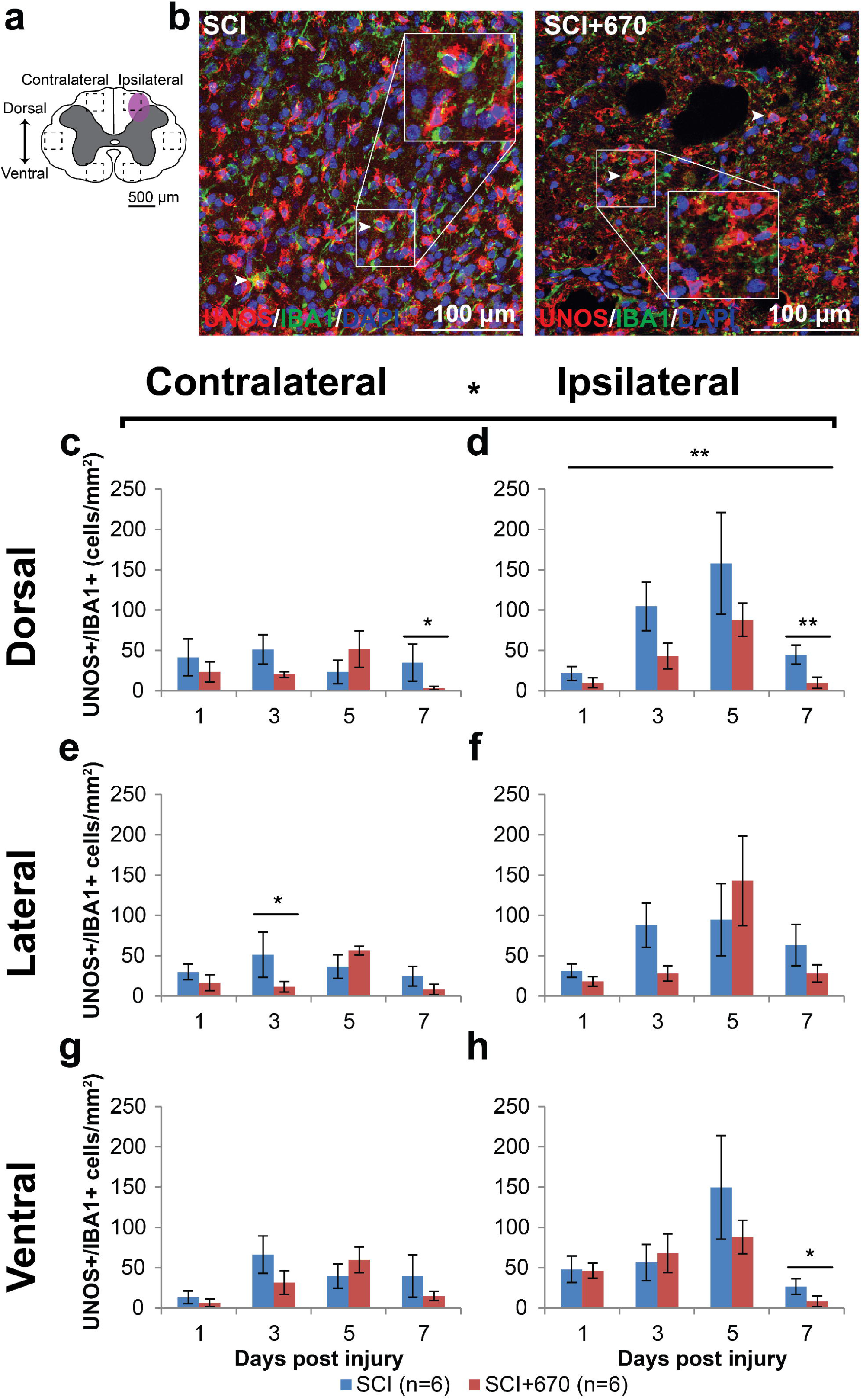
iNOS expressing microglia/macrophage immunoreactivity is reduced by red-light treatment following T10 hemi-contusion. **(a)** The schematic representation of the spinal cord illustrates the dorsal, lateral and ventral regions of interest for analysis (enclosed by dashed lines, area of each box: 0.1 mm^2^). Approximate location of injury is indicated by the purple shaded are. **(b)** Example images are shown of UNOS (red), IBA1 (green), and DAPI (blue) triple positive cells from spinal cord injured untreated and light-treated animals at the dorsal region, ipsilateral to the injury at 7-dpi. **(c-d)** Quantification of UNOS^+^IBA1^+^DAPI^+^ cells, expressed as triple positive cell density within the dorsal region of interest, is shown contralateral **(c)** and ipsilateral **(d)** to the injury of untreated and light-treated groups. **(e-f)** UNOS^+^IBA1^+^DAPI^+^ cell density in the lateral regions of interest contralateral **(e)** and ipsilateral **(f)** to the injury. **(g-h)** UNOS^+^IBA1^+^DAPI^+^ cell density in the ventral regions of interest contralateral **(g)** and ipsilateral **(h)** to the injury. n values indicated (legend) are for each time point. Data is expressed as mean ± SEM; * p < 0.05, ** p < 0.01, LMER.

In the dorsal region of the spinal cord, UNOS^+^IBA1^+^ cells were maintained below 50 per mm^2^ on the contralateral side throughout the recovery period in both groups **(Fig 9c).** However, on the ipsilateral side **(Fig 9d),** the 5-dpi time was significantly elevated compared to the 1- and 7-dpi times (p = 0.0010 and 0.0075 respectively, LMER, Tukey), and a similar pattern was evident in the ipsilateral ventral region **(Fig 9h)** between the 5- and 7-dpi times (p = 0.0003, LMER, Tukey).

While red-light significantly reduced UNOS^+^IBA1^+^ cell density overall, the effects arose mainly from the dorsal ipsilateral region across all time points **(Fig 9d),** and most strongly at the 7-dpi time.

These results show that iNOS expressing IBA1+ cells are present, predominantly on the ipsilateral side following hemi-contusion SCI, and red-light treatment reduces iNOS expression, particularly at the injury focus in the ipsilateral dorsal region.

## Discussion

Over the past decade, the use of photobiomodulation as a non-invasive therapy to improve repair of the injured nervous system and to reduce pain has gained increasing attention. Here, we characterized changes in mechanical sensitivity and spinal cord cellular environment in the subacute phase following a mild weight-drop hemi-contusion SCI and red-light (670 nm) treatment. We demonstrate that daily 30 min treatments of 670 nm at 35 mW/cm^2^ reduces the level of mechanical sensitivity in both SCI and sham-operated rats, as well as the overall incidence of animals developing hypersensitivity following SCI. These functional improvements in spinal cord injured animals were accompanied by reduced neuronal cell death, reduced iNOS expression in IBA1^+^ microglia/macrophages, and reduced astrocyte activation but not the IL1β^+^ astrocyte subpopulation in the injured spinal cord.

The current study used an LED with 35 W/cm^2^ to treat the animals over a transparent box. This brought the light source 10 mm away from direct contract to the skin surface. While placing the LED closer to the animal would increase irradiance, and therefore improve penetration of the red light, we have previously shown that 670 nm penetration is not linear to irradiance^32^ Photobiomodulation may involve both cellular or humoral factors^33^, yet the optimal and/or therapeutic window of irradiance for these differing mechanisms remains unknown. As such, dedicated studies would be required to determine the impact of removing the transparent box on animal recovery, as increasing light intensity can potentially be harmful^34^.

The treatment in the current study was carried out on animals with acute spinal cord injury. To date, complete studies examining the effects of photobiomodulation on chronic spinal cord injury remains limited, however a clinical trial is currently recruiting participants^35^ where investigators plan to implant an irradiation fibre into the injury site to treat acute spinal cord injury. Result for chronic brain injury have been encouraging; one study has shown that 18 sessions of LED treatment (22.2□mW/cm^2^) over 6 weeks significantly improved the cognition of patients with chronic mild traumatic brain injury^36^.

We used a mechanical testing paradigm that allowed assessment of above-, at-, and below-injury dermatomes, which provides an insight as to how mechanical sensitivity is changed following SCI. Categories I and II are not characteristic of nocifensive behaviours in laboratory rats while categories III and IV are inexorable behavioural responses to aversive stimuli including basic/integrated motor responses (jumping, avoidance and aggression) and vocalization^37^ As the CON group generally displayed categories I and II responses **(Fig 1a),** the mechanical sensitivity used in the current study does not induce pain in normal rats. The paradigm used to assess mechanical sensitivity in the present study presents some advantages over conventional assessment methods, such as using von Frey filaments to assess paw withdrawal threshold^20,38–40^, which requires complete intact motor function of the respective limbs^41^. In addition, the display of nocifensive behaviour is not affected by hind limb(s) motor deficits; animals are capable of avoidance behaviours using the front limbs only.

Using the pre-defined hypersensitivity threshold based on responses from normal intact animals, we were able to determine the percentage of animals developing hypersensitivity. This variation in the development of mechanical sensitivity is also observed in spinal cord injured patients^3,5,6^. SCI-induced mechanical allodynia was observed in at least 50% of the rats from 1-dpi, while the subpopulation developing hypersensitivity was reduced by 670 nm treatment from 1-dpi to 5-dpi **(Table 1)**. This mechanical hypersensitivity incidence in untreated spinal injured animals is consistent with another study which reported 67%^42^, and with our previous study of hypersensitivity incidence at 7-dpi using a more severe SCI model^21^. The current study provides the first evidence that 670 nm treatment reduces hypersensitivity incidence in the subacute phase following SCI in rats. There could be two possible explanations for the early reduction in hypersensitivity incidence; 670 nm light could either delay the development, or accelerate the recovery, of mechanical hypersensitivity. The latter is likely since 670 nm treatment has been shown to accelerate cellular processes and increase metabolism^43–45^. Furthermore, the peak hypersensitivity in the light-treated group was at 5-dpi **(Sup 2),** while the untreated injured group appeared to continue to develop hypersensitivity over the 7-day recovery period, and therefore, might require more time to fully develop maximal hypersensitivity.

As expected, only a few sham-injured animals were categorised as hypersensitive, suggesting that the surgical procedure does not produce excessive pain in the dorsum of the animal. Hypersensitive untreated spinal cord injured animals develop mechanical hypersensitivity mostly in the At-Level regions from 1-dpi with the potential to develop Below-Level hypersensitivity by 7-dpi. These results are consistent with spinal cord injured patients in clinical settings. Neuropathic pain patients develop both early-onset At-Level and late-onset Below-Level allodynia/hyperalgesia^5,7,8,31,46,47^. Although changes at the spinal and the supraspinal level have been correlated with the development of these two distinct pain phenotypes, the exact underlying mechanism remains unclear^5,48–50^. As we previously showed that the reduction in mechanical sensitivity is not due to reduced functional integrity of the dorsal column pathway^21^, it may arises from the anterolateral system. Following the surgery, bradykinins are released which elicits pain/sensitisation in the skin, while endogenous opioids such as endorphins are produced to control the pain^51^. Photobiomodulation has been shown to decrease bradykinins while increasing endorphins^52,53^, therefore the effect of red-light in the present study may result from alterations to bradykinins and/or endogenous opioids, leading to an overall reduction in sensitivity across all groups.

Little is known about the effect of red-light treatment on myelination. In the current study, no effect of 670 nm was observed on the level of myelination. The percentage area stained with MBP was found to be around 56% which is consistent with a previous study^54^. We and others show that demyelination starts within 24 h post SCI^55,56^. Remyelination following SCI starts around 7-dpi^56^, which is consistent with our MBP levels similar to CONs at 7-dpi **(Fig 3).** Up to date, there have been no studies documenting the effect of 670 nm treatment on demyelination following SCI. The most relevant study using 650 nm following sciatic nerve injury found no effect on number of Schwann cells in rabbits at 30 dpi^57^ It has been shown that demyelination also occurs in rostral and caudal regions^58^. Further investigations are thus needed to examine the effect of 670 nm on myelination outside the injury epicentre.

Studies have shown rapid loss of NF200 (especially dephosphorylated population) following SCI within the first 24 h^59^, which was consistent with our findings **(Fig 4).** Disrupted and degraded neurofilament then leads to axonal degeneration^60,61^, which could contribute to pain hypersensitivity. 670 nm treatment was not effective in stalling the loss of NF200. A similar result was found in a model of sciatic nerve injury in rabbits, where no effect of 650 nm was found on the number of myelinated axons at 30 dpi^57^ It is therefore worth investigating changes in axons rostral and caudal to the injury, as loss of neurofilament is well documented in those areas following SCI^58^

Neuronal cells undergo apoptosis and necrosis following SCI, while there is minimal neuronal death in naïve spinal cords^62–64^ Most neuronal death was observed in the dorsal and intermediate levels which would give rise to neurons in the spinothalamic tract, visceral motor neurons and interneurons. It is therefore not feasible to speculate which type(s) is more susceptible to cell death. 670 nm treatment has shown to significantly reduce neuronal cell death following SCI from 1-dpi. A similar study using 810 nm treatment showed 50% less motor neuron death following ischemia-induced SCI at 3-dpi^65^. The mechanism by which 670 nm treatment reduces neuronal cell death might be related to reduced iNOS **(Fig 9).** Xu and colleagues have shown that iNOS induces spinal neuronal degeneration following SCI through extracellular signal regulated kinases^66^.

Following SCI, pro-inflammatory cytokines are secreted by activated glial cells, both microglia/macrophages and astrocytes. During the post-injury period, these glial cells are activated through a combination of different mechanisms that involve neuroinflammation, ionic imbalance and cytokines/chemokines^46^. Regional and global increases in GFAP expression have been documented from 2 h persisting for at least 6 months^46^, in agreement with our observation of increased GFAP expression throughout the cord segment within 24 h post-injury, which persisted for at least 7 days **(Fig 6).** Limited studies have shown the effect of 670 nm treatment following SCI on astrocyte activation or GFAP upregulation, although similar effects using 810 nm have been documented^22^. Other groups have previously demonstrated a reduction in GFAP expression following light treatment in Müller cells in the retina^67,68^, and a decrease in GFAP expression by 670 nm irradiation in the brain of monkeys with Parkinson’s disease^69^. The present study is the first demonstration that GFAP upregulation in astrocytes is reduced by daily 670 nm treatment from 3-dpi following SCI. The reduced GFAP cannot account for the early (i.e. 1-dpi) red-light induced pain relief, as GFAP expression was not reduced at this time. Surprisingly, the reduction in GFAP did not appear to arise from the IL1β astrocyte subpopulation **(Fig 7).** IL1β is strongly implicated in the development of pain hypersensitivity^70^. Subpopulations of GFAP^+^ astrocytes have been reported before and IL1β^+^ astrocytes are considered to be pro-inflammatory and neurotoxic^71^. However, it is possible that red-light modulates other mediators or astrocyte subpopulations.

IL1β, produced by both astrocytes and microglia/macrophages, is a cytokine that not only augments inflammation, but also acts on both pre- and post-synaptic terminals to initiate and maintain pain^70,72^ We found no effect of 670 nm treatment on IL1β production in microglia/macrophages following SCI from 1 to 7-dpi **(Fig 8).** This observation is consistent with our previous study^21^ showing no effect of 670 nm on pro-inflammatory microglia/macrophage (M1), and another study where the authors showed no effect of 810 nm on IL1β expression at either 6 h post injury or 4-dpi following SCI^22^. However, other studies have reported decreases in IL1 β expression by light treatment in other injury models^73,74^ This discrepancy suggests that light treatment may activate different cell pathways in different injury models. As IL1β is a significant player in the regulation of pain, the present study suggests that the pain-alleviating effect of 670 nm light treatment may be independent, or downstream to IL1β, at least up to the first 7 days. Furthermore, we found a reduction in the behavioural sensitivity of red-light treated sham-injured animals **(Fig 1c).** The mechanism of this red-light-induced reduction of sensitivity in these animals is also unlikely to be IL1 β-dependant because these animals displayed no evidence of pain behaviours.

We also found a significant decrease in UNOS expression, a surrogate marker for iNOS in microglia/macrophages, following 670 nm treatment in spinal cord injured animals **(Fig 9).** Byrnes *et al.* found a significant reduction in iNOS transcription that was measured by RT-PCR at 6 h post-injury, but not at 4-dpi using 810 nm treatment^22^ The discrepancy between our study and that of Byrnes *et al.* at the 3 to 4-dpi might indicate a different mechanism of action between 670 nm and 810 nm, or that the reduction of iNOS^+^ microglia/macrophages was masked by an increase of iNOS in other cells types not observed by Byrnes *et al*. It is interesting that IL1β and iNOS, both considered markers of classically activated microglia/macrophages (M1), were affected differently by red-light treatment. In our previous study, we demonstrated that red-light did not alter the proportion of M1 cells, however, the proportion of M2 cells was increased^21^. The current study is partly consistent with this idea; IL1 β expressing cells (M1 phenotype) were not altered by red-light treatment, however immunoreactive iNOS (another marker of M1 phenotype) microglia/macrophages were reduced. iNOS down-regulation may indirectly indicate conversion towards arginase upregulation^75^ and therefore conversion towards the M2 phenotype. It may be possible that iNOS and IL1β are produced by different cells and that the decrease in iNOS expression in microglia/macrophages might occur in response to the increase in M2 microglia/macrophage^21^. In addition to the beneficial effect of a less inflammatory microenvironment, the decreased iNOS expression in microglia/macrophages could also contribute to the reduced mechanical sensitivity in 670 nm treated animals. Studies have shown that pain following SCI could be reduced, or reversed, by iNOS inhibitors or general NOS inhibition^76,77^ The present study supports the idea that 670 nm may act on a pain modulation pathway that is iNOS-dependent, but IL1 β-independent. Future studies examining these ideas would be of interest. Moreover, future studies using stereology, such as the optical fractionator technique^78^, to quantify 670 nm-induced changes in neuron and glial cell numbers in the entire spinal cord structure would be of interest.

In conclusion, we demonstrate that daily 670 nm irradiation reduces mechanical sensitivity in spinal cord injured and in sham operated rats, and reduces the chance of developing hypersensitivity up to 5-days post SCI. The analgesic effects of red-light therapy in the subacute stage after SCI may involve reduced neuronal death, reduced astrogliosis and reduced iNOS expression in microglia/macrophage in the spinal cord, independent of myelination and IL1β glial cell expression. The combined reduction of iNOS^+^ microglia/macrophages and IL1β^-^ astrocytes may therefore impact different stages of mechanical sensitivity following SCI. Taken together, these findings suggest that red-light therapy can be a safe non-pharmacological approach to manage pain by modulating the endogenous response of neuronal and glial populations following SCI.

## Acknowledgements

The authors gratefully acknowledge the Bootes Medical Research Foundation for funding this project.

---

**Sup 1.**
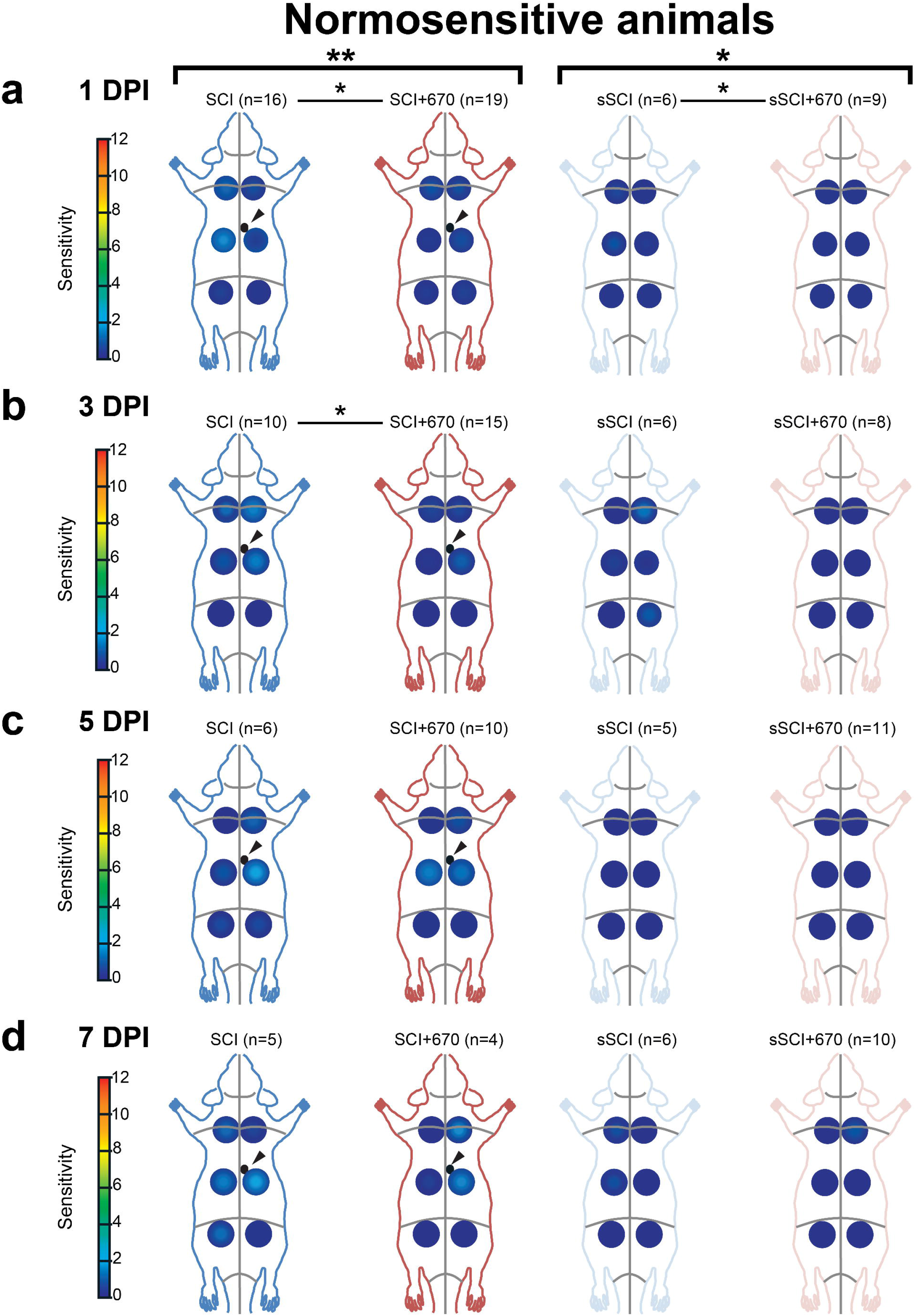
Red-light treatment reduces regional sensitivity scores (RSSs) in normosensitive animals. RSSs in SCI (blue), SCI+670 (red), sSCI (light blue), and sSCI+670 (pink) animals at **(a)** 1-dpi, **(b)** 3-dpi, **(c)** 5-dpi, and **(d)** 7-dpi are shown for animals that did not develop hypersensitivity. Arrowheads indicate location of T10 hemi-contusion injury (small back circles) in spinal cord injured groups. Statistical comparisons between 2 groups across all time points (black bracket, CLMM), across all levels at individual time point (black lines, CLMM) and between 2 groups at different levels (grey lines, CLMM) are indicated. Data is expressed as mean ± SEM as per **Fig 1a** inset; n values indicated for each group; * p < 0.05, ** p < 0.01, CLMM. See **Fig 1** for abbreviations.

**Sup 2.**
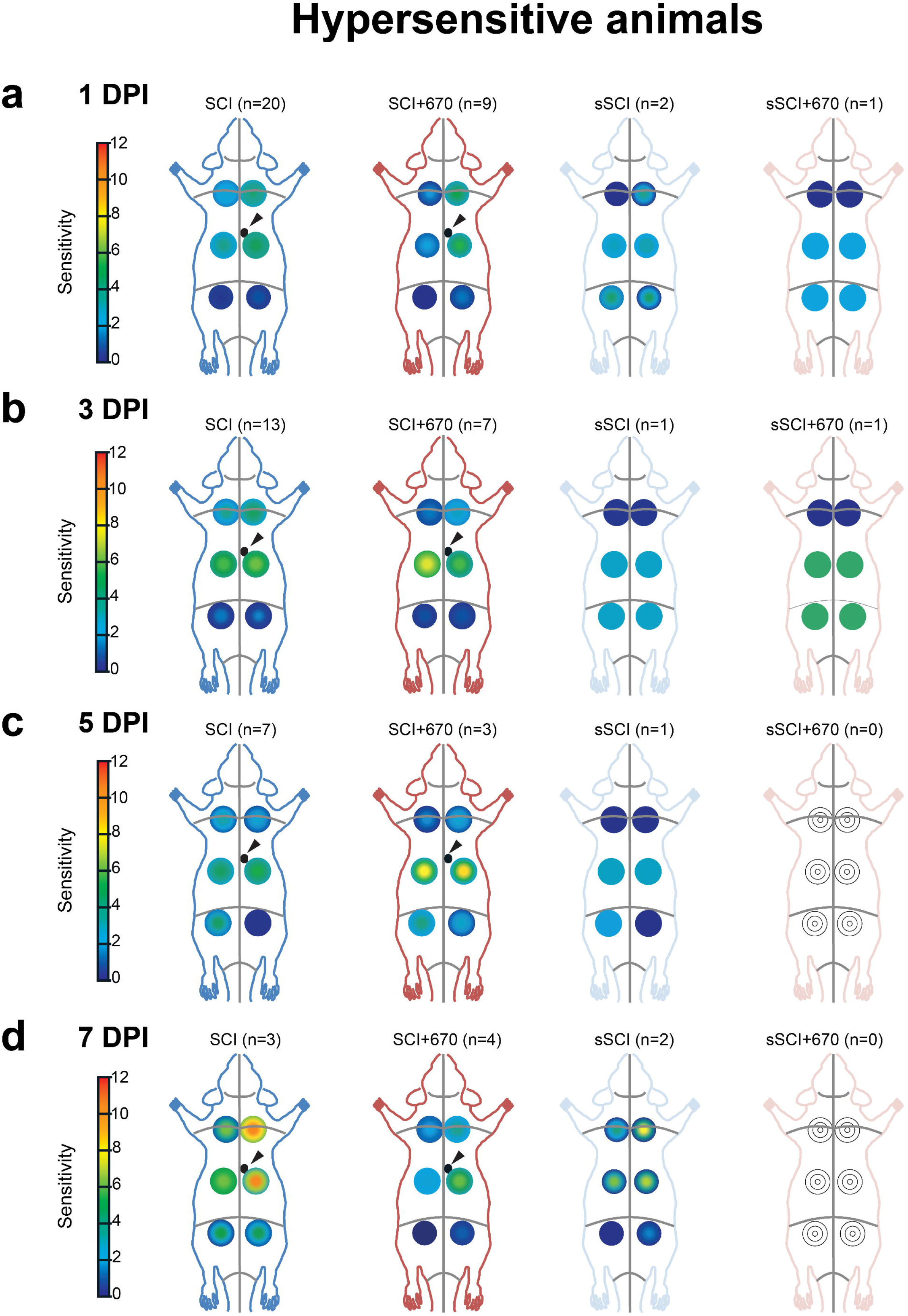
Regional sensitivity scores (RSSs) in hypersensitive animals. RSSs in SCI (blue), SCI+670 (red), sSCI (light blue), and sSCI+670 (pink) animals at **(a)** 1-dpi, **(b)** 3-dpi, **(c)** 5-dpi, and **(d)** 7-dpi are shown for animals that developed hypersensitivity. Arrowheads indicate location of T10 hemi-contusion injury (small back circles) in spinal cord injured groups. Statistical comparisons between 2 groups across all time points (black bracket, CLMM), across all levels at individual time point (black lines, CLMM) and between 2 groups at different levels (grey lines, CLMM) are indicated. Data is expressed as mean ± SEM as per **Fig 1a** inset; n values indicated for each group. See **Fig 1** for abbreviations.

**Sup 3.**
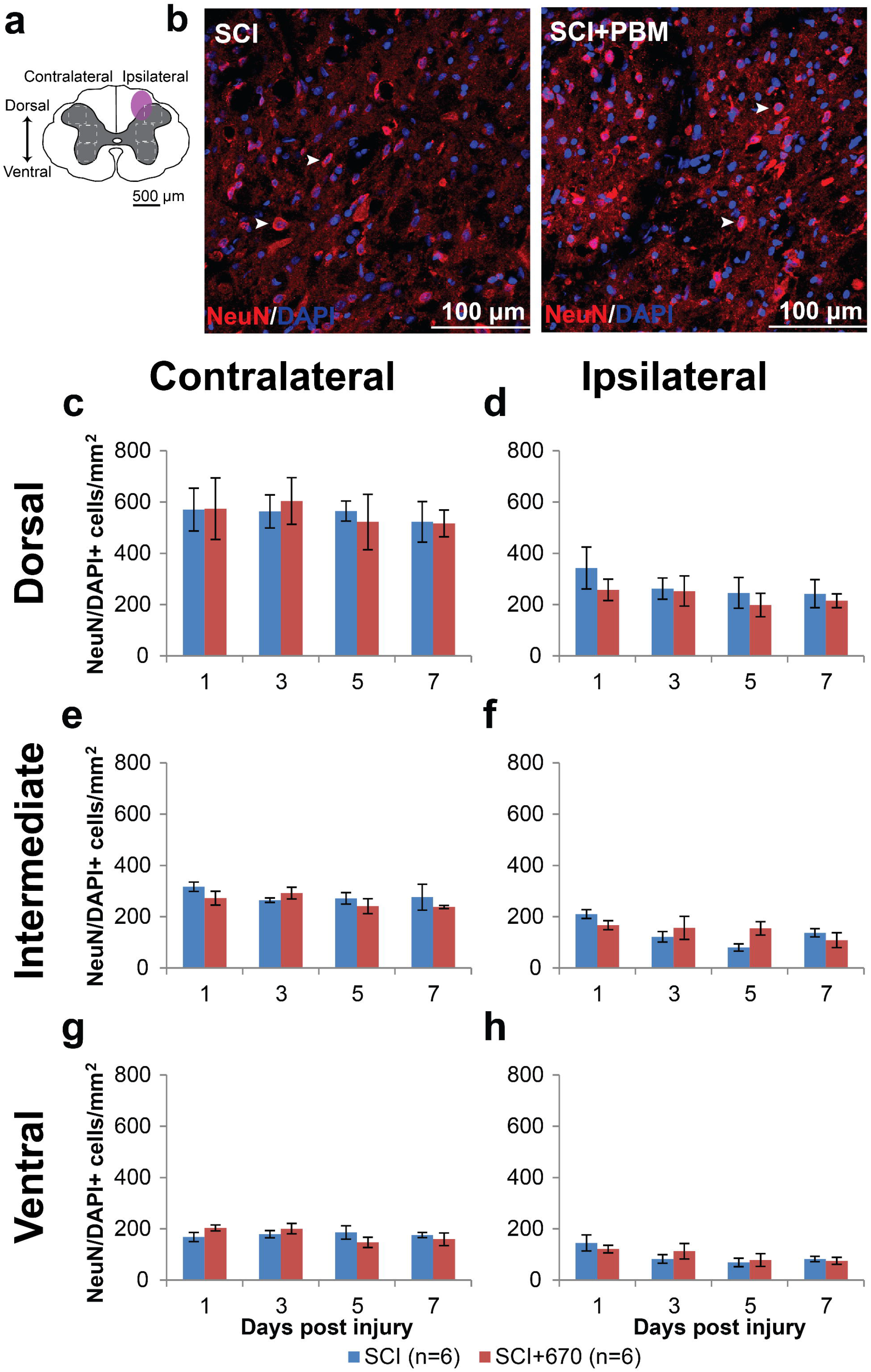
Neuronal cell density is reduced ipsilateral to SCI but not altered by 670 nm treatment. **(a)** The schematic representation of the spinal cord illustrates the dorsal, intermediate and ventral regions of interest for analysis (enclosed by dashed lines, 0.1 mm^2^). Approximate location of injury epicentre is indicated by the purple shaded area. **(b)** Example images of NeuN (red) and DAPI (blue) double positive cells from spinal cord injured untreated and light-treated groups ipsilateral to the injury at dorsal region at 1-dpi. **(c-d)** Quantification of NeuN^+^DAPI^+^ cells, expressed as double positive cell density within the region of interest, in the dorsal region of the spinal cord, contralateral **(c)** and ipsilateral **(d)** to the injury of untreated and light-treated groups. **(e-f)** NeuN^+^DAPI^+^ cell density in the intermediate regions of interest contralateral **(e)** and ipsilateral **(f)** to the injury. **(g-h)** NeuN^+^DAPI^+^ cell density in the ventral regions of interest contralateral (g) and ipsilateral (h) to the injury. Data is expressed as mean ± SEM; n values indicated (legend) are for each time point.

**Sup 4.**
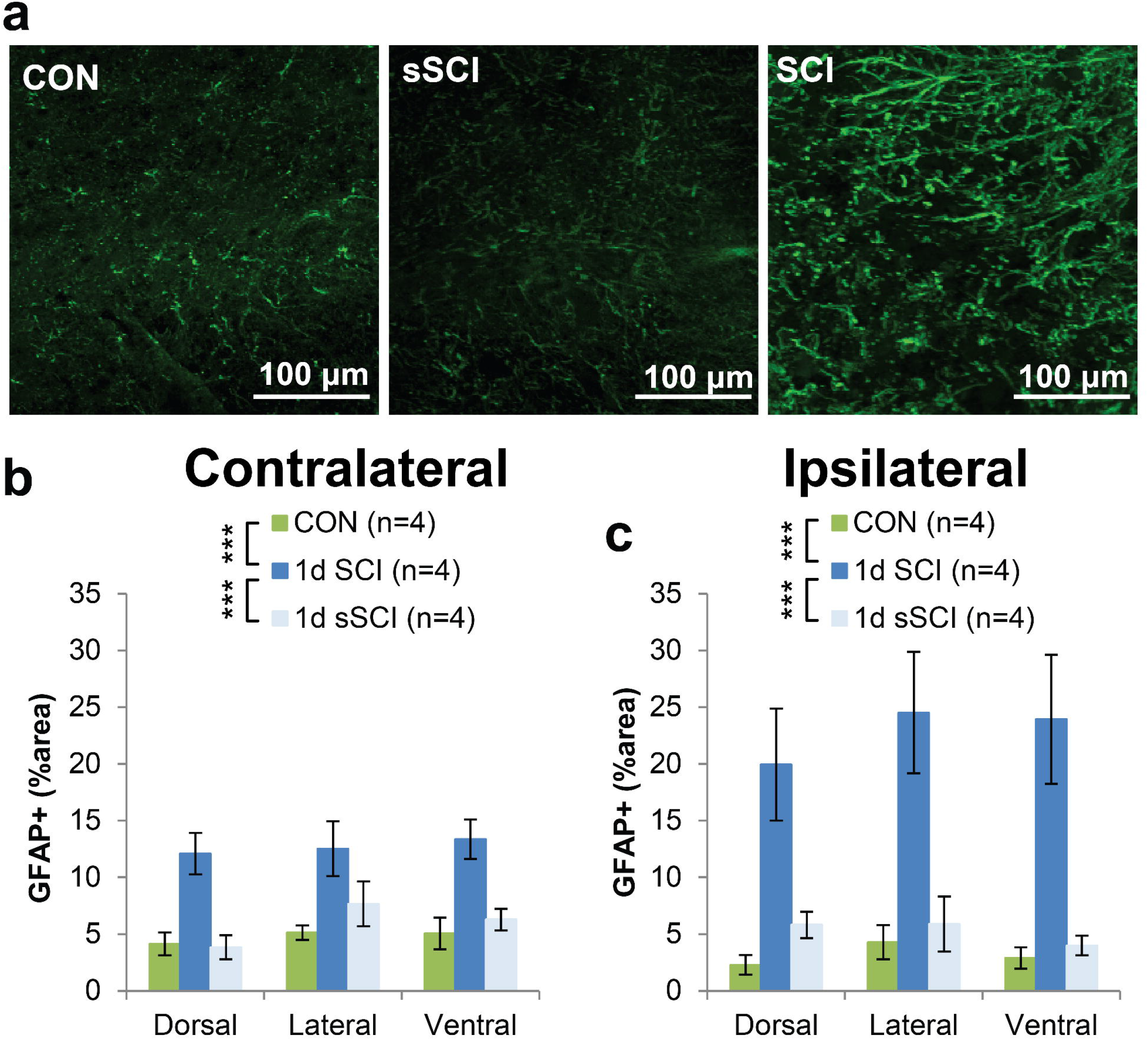
GFAP increases within 24 h following a mild T10 hemi-contusion injury. **(a)** Example images of GFAP^+^ staining (green) from uninjured, spinal cord injured (1-dpi) and sham-injured (1-dpi) animals ipsilateral to the injury at the dorsal level. **(b-c)** Quantification of GFAP^+^ labelling at 1 day postinjury, expressed as the percentage area of positive label above threshold within dorsal, lateral and ventral regions of interest (see **Fig 6A),** contralateral **(b)** and ipsilateral **(c)** to the injury of uninjured controls, spinal cord injured (1-dpi) and sham-injured (1-dpi) groups. Vertical brackets indicate statistical comparisons among three groups across all regions (LMER). Data is expressed as mean ± SEM; *** p < 0.001. See **Fig 1** for abbreviations.

## References

1. World Health Organization and International Spinal Cord Society (2013). International perspectives on spinal cord injury. World Health Organization.

2. Singh, A., Tetreault, L., Kalsi-Ryan, S., Nouri, A. and Fehlings, M.G. (2014). Global prevalence and incidence of traumatic spinal cord injury. Clin Epidemiol 6, 309–331.

3. Turner, J.A., Cardenas, D.D., Warms, C.A. and McClellan, C.B. (2001). Chronic pain associated with spinal cord injuries: a community survey. Arch Phys Med Rehabil 82, 501–509.

4. Pascoal-Faria, P., Yalcin, N. and Fregni, F. (2015). Neural markers of neuropathic pain associated with maladaptive plasticity in spinal cord injury. Pain Pract 15, 371–377.

5. Finnerup, N.B., Norrbrink, C., Trok, K., Piehl, F., Johannesen, I.L., Sorensen, J.C., Jensen, T.S. and Werhagen, L. (2014). Phenotypes and predictors of pain following traumatic spinal cord injury: a prospective study. J Pain 15, 40–48.

6. Dijkers, M., Bryce, T. and Zanca, J. (2009). Prevalence of chronic pain after traumatic spinal cord injury: a systematic review. J Rehabil Res Dev 46, 13–29.

7. Bryce, T.N. (2009). Spinal cord injury. Springer Publishing Company.

8. Widerstrom-Noga, E., Biering-Sorensen, E., Bryce, T.N., Cardenas, D.D., Finnerup, N.B., Jensen, M.P., Richards, J.S. and Siddall, P.J. (2014). The International Spinal Cord Injury Pain Basic Data Set (version 2.0). Spinal Cord 52, 282–286.

9. Finnerup, N.B., Johannesen, I.L., Fuglsang-Frederiksen, A., Bach, F.W. and Jensen, T.S. (2003). Sensory function in spinal cord injury patients with and without central pain. Brain 126, 57–70.

10. D’Angelo, R., Morreale, A., Donadio, V., Boriani, S., Maraldi, N., Plazzi, G. and Liguori, R. (2013). Neuropathic pain following spinal cord injury: what we know about mechanisms, assessment and management. Eur Rev Med Pharmacol Sci 17, 3257–3261.

11. Chiang, C.Y., Sessle, B.J. and Dostrovsky, J.O. (2012). Role of astrocytes in pain. Neurochem Res 37, 2419–2431.

12. Nakajima, A., Tsuboi, Y., Suzuki, I., Honda, K., Shinoda, M., Kondo, M., Matsuura, S., Shibuta, K., Yasuda, M., Shimizu, N. and Iwata, K. (2011). PKCgamma in Vc and C1/C2 is involved in trigeminal neuropathic pain. J Dent Res 90, 777–781.

13. Kobayashi, A., Shinoda, M., Sessle, B.J., Honda, K., Imamura, Y., Hitomi, S., Tsuboi, Y., Okada-Ogawa, A. and Iwata, K. (2011). Mechanisms involved in extraterritorial facial pain following cervical spinal nerve injury in rats. Mol Pain 7, 12.

14. Ji, R.R., Gereau, R.W.t., Malcangio, M. and Strichartz, G.R. (2009). MAP kinase and pain. Brain Res Rev 60, 135–148.

15. Nakagawa, T. and Kaneko, S. (2010). Spinal astrocytes as therapeutic targets for pathological pain. J Pharmacol Sci 114, 347–353.

16. McMahon, S.B. and Malcangio, M. (2009). Current challenges in glia-pain biology. Neuron 64, 46–54.

17. Ikeda, H., Stark, J., Fischer, H., Wagner, M., Drdla, R., Jager, T. and Sandkuhler, J. (2006). Synaptic amplifier of inflammatory pain in the spinal dorsal horn. Science 312, 1659–1662.

18. Galan-Arriero, I., Avila-Martin, G., Ferrer-Donato, A., Gomez-Soriano, J., Bravo-Esteban, E. and Taylor, J. (2014). Oral administration of the p38alpha MAPK inhibitor, UR13870, inhibits affective pain behavior after spinal cord injury. Pain 155, 2188–2198.

19. Watanabe, S., Uchida, K., Nakajima, H., Matsuo, H., Sugita, D., Yoshida, A., Honjoh, K., Johnson, W.E. and Baba, H. (2015). Early transplantation of mesenchymal stem cells after spinal cord injury relieves pain hypersensitivity through suppression of pain-related signaling cascades and reduced inflammatory cell recruitment. Stem Cells 33, 1902–1914.

20. Tateda, S., Kanno, H., Ozawa, H., Sekiguchi, A., Yahata, K., Yamaya, S. and Itoi, E. (2017). Rapamycin suppresses microglial activation and reduces the development of neuropathic pain after spinal cord injury. J Orthop Res 35, 93–103.

21. Hu, D., Zhu, S. and Potas, J.R. (2016). Red LED photobiomodulation reduces pain hypersensitivity and improves sensorimotor function following mild T10 hemicontusion spinal cord injury. J Neuroinflammation 13, 200.

22. Byrnes, K.R., Waynant, R.W., Ilev, I.K., Wu, X., Barna, L., Smith, K., Heckert, R., Gerst, H. and Anders, J.J. (2005). Light promotes regeneration and functional recovery and alters the immune response after spinal cord injury. Lasers in surgery and medicine 36, 171–185.

23. Yang, X., Askarova, S., Sheng, W., Chen, J.K., Sun, A.Y., Sun, G.Y., Yao, G. and Lee, J.C. (2010). Low energy laser light (632.8 nm) suppresses amyloid-beta peptide-induced oxidative and inflammatory responses in astrocytes. Neuroscience 171, 859–868.

24. Vijayaprakash, K.M. and Sridharan, N. (2013). An experimental spinal cord injury rat model using customized impact device: A cost-effective approach. Journal of pharmacology & pharmacotherapeutics 4, 211–213.

25. Schneider, C.A., Rasband, W.S. and Eliceiri, K.W. (2012). NIH Image to ImageJ: 25 years of image analysis. Nat Methods 9, 671–675.

26. R Core Team (2016). R: A language and environment for statistical computing. R Foundation for Statistical Computing: Vienna, Austria.

27. Kuznetsova, A., Brockhoff, P.B. and Christensen, R.H.B. (2016). lmerTest: Tests in Linear Mixed Effects Models.

28. McElreath, R. (2016). AIC provides a surprisingly simple estimate of the average out-of-sample deviance. In: Statistical Rethinking: A Bayesian Course with Examples in R and Stan CRC Press, pps. 189.

29. Christensen, R.H.B. (2015). Package ‘ordinal’.

30. Ji, R.R., Berta, T. and Nedergaard, M. (2013). Glia and pain: is chronic pain a gliopathy? Pain 154 Suppl 1, S10–28.

31. Hulsebosch, C.E., Hains, B.C., Crown, E.D. and Carlton, S.M. (2009). Mechanisms of chronic central neuropathic pain after spinal cord injury. Brain Res Rev 60, 202–213.

32. Hu, D., van Zeyl, M., Valter, K. and Potas, J.R. (2019). Sex, but not skin tone affects penetration of red-light (660 nm) through sites susceptible to sports injury in lean live and cadaveric tissues. Journal of Biophotonics 0, e201900010.

33. Johnstone, D.M., Moro, C., Stone, J., Benabid, A.L. and Mitrofanis, J. (2015). Turning On Lights to Stop Neurodegeneration: The Potential of Near Infrared Light Therapy in Alzheimer’s and Parkinson’s Disease. Front Neurosci 9, 500.

34. Chu-Tan, J.A., Rutar, M., Saxena, K., Wu, Y., Howitt, L., Valter, K., Provis, J. and Natoli, R. (2016). Efficacy of 670 □nm Light Therapy to Protect against Photoreceptor Cell Death Is Dependent on the Severity of Damage. International Journal of Photoenergy 2016, 1–12.

35. Hospital, X. (2018). Clinical Study of Treatment of Acute Spinal Cord Injury by Near Infrared Light Irradiation. https://ClinicalTrials.gov/show/NCT03643419.

36. Naeser, M.A., Zafonte, R., Krengel, M.H., Martin, P.I., Frazier, J., Hamblin, M.R., Knight, J.A., Meehan, W.P., 3rd and Baker, E.H. (2014). Significant improvements in cognitive performance post-transcranial, red/near-infrared light-emitting diode treatments in chronic, mild traumatic brain injury: open-protocol study. J Neurotrauma 31, 1008–1017.

37. Institute for Laboratory Animal Research (U.S.). Committee on Recognition and Alleviation of Pain in Laboratory Animals. (2009). Recognition and alleviation of pain in laboratory animals. National Academies Press: Washington, D.C.

38. Kanno, H., Pressman, Y., Moody, A., Berg, R., Muir, E.M., Rogers, J.H., Ozawa, H., Itoi, E., Pearse, D.D. and Bunge, M.B. (2014). Combination of engineered Schwann cell grafts to secrete neurotrophin and chondroitinase promotes axonal regeneration and locomotion after spinal cord injury. The Journal of neuroscience: the official journal of the Society for Neuroscience 34, 1838–1855.

39. Bartus, K., James, N.D., Didangelos, A., Bosch, K.D., Verhaagen, J., Yanez-Munoz, R.J., Rogers, J.H., Schneider, B.L., Muir, E.M. and Bradbury, E.J. (2014). Large-scale chondroitin sulfate proteoglycan digestion with chondroitinase gene therapy leads to reduced pathology and modulates macrophage phenotype following spinal cord contusion injury. The Journal of neuroscience: the official journal of the Society for Neuroscience 34, 4822–4836.

40. Yu, D., Thakor, D.K., Han, I., Ropper, A.E., Haragopal, H., Sidman, R.L., Zafonte, R., Schachter, S.C. and Teng, Y.D. (2013). Alleviation of chronic pain following rat spinal cord compression injury with multimodal actions of huperzine A. Proc Natl Acad Sci U SA 110, E746–755.

41. Detloff, M.R., Clark, L.M., Hutchinson, K.J., Kloos, A.D., Fisher, L.C. and Basso, D.M. (2010). Validity of acute and chronic tactile sensory testing after spinal cord injury in rats. Exp Neurol 225, 366–376.

42. Siddall, P., Xu, C.L. and Cousins, M. (1995). Allodynia following traumatic spinal cord injury in the rat. Neuroreport 6, 1241–1244.

43. Begum, R., Calaza, K., Kam, J.H., Salt, T.E., Hogg, C. and Jeffery, G. (2015). Near-infrared light increases ATP, extends lifespan and improves mobility in aged Drosophila melanogaster. Biol Lett 11.

44. Yip, K.K., Lo, S.C., Leung, M.C., So, K.F., Tang, C.Y. and Poon, D.M. (2011). The effect of low-energy laser irradiation on apoptotic factors following experimentally induced transient cerebral ischemia. Neuroscience 190, 301–306.

45. Fells, J.T., Henry, M.M., Summerfelt, P., Wong-Riley, M.T., Buchmann, E.V., Kane, M., Whelan, N.T. and Whelan, H.T. (2003). Therapeutic photobiomodulation for methanol-induced retinal toxicity. Proc Natl Acad Sci USA 100, 3439–3444.

46. Gwak, Y.S., Kang, J., Unabia, G.C. and Hulsebosch, C.E. (2012). Spatial and temporal activation of spinal glial cells: role of gliopathy in central neuropathic pain following spinal cord injury in rats. Exp Neurol 234, 362–372.

47. Finnerup, N.B., Baastrup, C. and Jensen, T.S. (2009). Neuropathic pain following spinal cord injury pain: mechanisms and treatment. Scandinavian Journal of Pain 1, S3–S11.

48. Siddall, P.J., McClelland, J.M., Rutkowski, S.B. and Cousins, M.J. (2003). A longitudinal study of the prevalence and characteristics of pain in the first 5 years following spinal cord injury. Pain 103, 249–257.

49. Wrigley, P.J., Press, S.R., Gustin, S.M., Macefield, V.G., Gandevia, S.C., Cousins, M.J., Middleton, J.W., Henderson, L.A. and Siddall, P.J. (2009). Neuropathic pain and primary somatosensory cortex reorganization following spinal cord injury. Pain 141, 52–59.

50. Hari, A.R., Wydenkeller, S., Dokladal, P. and Halder, P. (2009). Enhanced recovery of human spinothalamic function is associated with central neuropathic pain after SCI. Exp Neurol 216, 428–430.

51. Stein, C., Clark, J.D., Oh, U., Vasko, M.R., Wilcox, G.L., Overland, A.C., Vanderah, T.W. and Spencer, R.H. (2009). Peripheral mechanisms of pain and analgesia. Brain Res Rev 60, 90–113.

52. Lins, R.D., Dantas, E.M., Lucena, K.C., Catao, M.H., Granville-Garcia, A.F. and Carvalho Neto, L.G. (2010). Biostimulation effects of low-power laser in the repair process. An Bras Dermatol 85, 849–855.

53. Chaves, M.E., Araujo, A.R., Piancastelli, A.C. and Pinotti, M. (2014). Effects of low-power light therapy on wound healing: LASER x LED. An Bras Dermatol 89, 616–623.

54. Crocker, S.J., Whitmire, J.K., Frausto, R.F., Chertboonmuang, P., Soloway, P.D., Whitton, J.L. and Campbell, I.L. (2006). Persistent macrophage/microglial activation and myelin disruption after experimental autoimmune encephalomyelitis in tissue inhibitor of metalloproteinase-1-deficient mice. Am J Pathol 169, 2104–2116.

55. Lasiene, J., Shupe, L., Perlmutter, S. and Horner, P. (2008). No evidence for chronic demyelination in spared axons after spinal cord injury in a mouse. The Journal of neuroscience: the official journal of the Society for Neuroscience 28, 3887–3896.

56. Totoiu, M.O. and Keirstead, H.S. (2005). Spinal cord injury is accompanied by chronic progressive demyelination. J Comp Neurol 486, 373–383.

57. Takhtfooladi, M.A. and Sharifi, D. (2015). A comparative study of red and blue light-emitting diodes and low-level laser in regeneration of the transected sciatic nerve after an end to end neurorrhaphy in rabbits. Lasers Med Sci 30, 2319–2324.

58. Kozlowski, P., Raj, D., Liu, J., Lam, C., Yung, A.C. and Tetzlaff, W. (2008). Characterizing white matter damage in rat spinal cord with quantitative MRI and histology. J Neurotrauma 25, 653–676.

59. Schumacher, P.A., Eubanks, J.H. and Fehlings, M.G. (1999). Increased calpain I-mediated proteolysis, and preferential loss of dephosphorylated NF200, following traumatic spinal cord injury. Neuroscience 91, 733–744.

60. Park, E., Velumian, A.A. and Fehlings, M.G. (2004). The role of excitotoxicity in secondary mechanisms of spinal cord injury: a review with an emphasis on the implications for white matter degeneration. J Neurotrauma 21, 754–774.

61. Eftekharpour, E., Karimi-Abdolrezaee, S. and Fehlings, M.G. (2008). Current status of experimental cell replacement approaches to spinal cord injury. Neurosurg Focus 24, E19.

62. Rosado, I.R., Lavor, M.S., Alves, E.G., Fukushima, F.B., Oliveira, K.M., Silva, C.M., Caldeira, F.M., Costa, P.M. and Melo, E.G. (2014). Effects of methylprednisolone, dantrolene, and their combination on experimental spinal cord injury. Int J Clin Exp Pathol 7, 4617–4626.

63. Li, H.T., Zhao, X.Z., Zhang, X.R., Li, G., Jia, Z.Q., Sun, P., Wang, J.Q., Fan, Z.K. and Lv, G. (2016). Exendin-4 Enhances Motor Function Recovery via Promotion of Autophagy and Inhibition of Neuronal Apoptosis After Spinal Cord Injury in Rats. Mol Neurobiol 53, 4073–4082.

64. Zhang, Z., Huang, Z., Dai, H., Wei, L., Sun, S. and Gao, F. (2015). Therapeutic Efficacy of E-64-d, a Selective Calpain Inhibitor, in Experimental Acute Spinal Cord Injury. Biomed Res Int 2015, 134242.

65. Sotoudeh, A., Jahanshahi, A., Zareiy, S., Darvishi, M., Roodbari, N. and Bazzazan, A. (2015). The influence of low-level laser irradiation on spinal cord injuries following ischemia-reperfusion in rats. Acta Cir Bras 30, 611–616.

66. Xu, Z., Wang, B.R., Wang, X., Kuang F., Duan, X.L., Jiao, X.Y. and Ju, G. (2006). ERK1/2 and p38 mitogen-activated protein kinase mediate iNOS-induced spinal neuron degeneration after acute traumatic spinal cord injury. Life Sci 79, 1895–1905.

67. Begum, R., Powner, M.B., Hudson, N., Hogg, C. and Jeffery, G. (2013). Treatment with 670 nm light up regulates cytochrome C oxidase expression and reduces inflammation in an age-related macular degeneration model. PLoS One 8, e57828.

68. Marco, F.D., Romeo, S., Nandasena, C., Purushothuman, S., Adams, C., Bisti, S. and Stone, J. (2013). The time course of action of two neuroprotectants, dietary saffron and photobiomodulation, assessed in the rat retina. Am J Neurodegener Dis 2, 208–220.

69. El Massri, N., Moro, C., Torres, N., Darlot, F., Agay, D., Chabrol, C., Johnstone, D.M., Stone, J., Benabid, A.L. and Mitrofanis, J. (2016). Near-infrared light treatment reduces astrogliosis in MPTP-treated monkeys. Exp Brain Res 234, 3225–3232.

70. Calvo, M., Dawes, J.M. and Bennett, D.L. (2012). The role of the immune system in the generation of neuropathic pain. Lancet Neurol 11, 629–642.

71. Liddelow, S.A., Guttenplan, K.A., Clarke, L.E., Bennett, F.C., Bohlen, C.J., Schirmer, L., Bennett, M.L., Munch, A.E., Chung, W.S., Peterson, T.C., Wilton, D.K., Frouin, A., Napier, B.A., Panicker, N., Kumar, M., Buckwalter, M.S., Rowitch, D.H., Dawson, V.L., Dawson, T.M., Stevens, B. and Barres, B.A. (2017). Neurotoxic reactive astrocytes are induced by activated microglia. Nature 541, 481–487.

72. Clark, A.K., Gruber-Schoffnegger, D., Drdla-Schutting, R., Gerhold, K.J., Malcangio, M. and Sandkuhler, J. (2015). Selective activation of microglia facilitates synaptic strength. The Journal of neuroscience: the official journal of the Society for Neuroscience 35, 4552–4570.

73. Sahu, K., Sharma, M. and Gupta, P.K. (2015). Modulation of inflammatory response of wounds by antimicrobial photodynamic therapy. Laser Ther 24, 201–208.

74. Lee, W.J., Lee, K.C., Kim, M.J., Jang, Y.H., Lee, S.J. and Kim, D.W. (2016). Efficacy of Red or Infrared Light-Emitting Diodes in a Mouse Model of Propionibacterium acnes-Induced Inflammation. Ann Dermatol 28, 186–191.

75. Lee, J., Ryu, H., Ferrante, R.J., Morris, S.M., Jr. and Ratan, R.R. (2003). Translational control of inducible nitric oxide synthase expression by arginine can explain the arginine paradox. Proc Natl Acad Sci USA 100, 4843–4848.

76. Yu, Y., Matsuyama, Y., Nakashima, S., Yanase, M., Kiuchi, K. and Ishiguro, N. (2004). Effects of MPSS and a potent iNOS inhibitor on traumatic spinal cord injury. Neuroreport 15, 2103–2107.

77. Fairbanks, C.A., Schreiber, K.L., Brewer, K.L., Yu, C.G., Stone, L.S., Kitto, K.F., Nguyen, H.O., Grocholski, B.M., Shoeman, D.W., Kehl, L.J., Regunathan, S., Reis, D.J., Yezierski, R.P. and Wilcox, G.L. (2000). Agmatine reverses pain induced by inflammation, neuropathy, and spinal cord injury. Proc Natl Acad Sci USA 97, 10584–10589.

78. Olesen, M.V., Needham, E.K and Pakkenberg, B. (2017). The Optical Fractionator Technique to Estimate Cell Numbers in a Rat Model of Electroconvulsive Therapy. J Vis Exp.

